# An extended β-finger motif is near-ubiquitous in kinase fusion proteins

**DOI:** 10.1101/2025.07.14.664643

**Authors:** Oliver R. Powell, David Gilbert, Brande B. H. Wulff

## Abstract

Kinase fusion proteins (KFPs) are immune receptors that confer disease resistance in wheat (*Triticum aestivum*) and barley (*Hordeum vulgare*). We discovered that KFPs with a confirmed function share an extended β-finger motif, a Poaceae-specific feature that originated ∼98 million years ago. The genes encoding these receptors are among the most highly expressed within the plant KFP repertoire, suggesting that elevated transcript abundance is a hallmark of their disease resistance function.

## Main Text

Kinase fusion proteins (KFPs) are a diverse set of immune receptors that confer resistance to a range of pathogens in wheat (*Triticum aestivum*) and barley (*Hordeum vulgare*). Structural analysis of the KFPs, Sr62^TK^ and Rwt4, previously identified an unusual extended β-finger motif in at least one kinase domain of each protein. This motif participates in the recognition of the corresponding pathogen effectors AvrSr62 and Pwt4, respectively^1–3^. The motif also maintains Sr62^TK^ as a homodimer^3^. AlphaFold3 predictions of the 18 experimentally validated KFPs revealed that the majority contained at least one kinase domain with an extended β-finger motif (Fig S1-20, Table S1-2). Given the prevalence of the extended β-finger motif in KFPs with a validated disease resistance function, we speculated that this motif would be suitable for the *ab initio* identification of KFPs.

To investigate the extent of sequence conservation and diversity of the extended β-finger motif within KFPs, we extracted the sequences of the motif from the AlphaFold 3 structures (Table S3). Yr15 was excluded, as the β-finger motifs in its two kinase domains lack the extended conformation (Fig. S3). We then constructed an initial hidden Markov model (HMM) of the motif from an alignment of these sequences. We iteratively refined the HMM using the proteome of bread wheat cv. Chinese Spring to improve its specificity and sensitivity (Fig. S21)^4^.

This approach yielded a motif with a highly conserved length, although the amino acid sequences of the motif varied considerably across KFPs. Based on these structural predictions and sequences, we defined the extended β-finger as a 31-amino acid motif that extends the highly conserved 4^th^ and 5^th^ β-sheets of the kinase domain (Fig. 1a, b)^5^.

**Fig 1.**
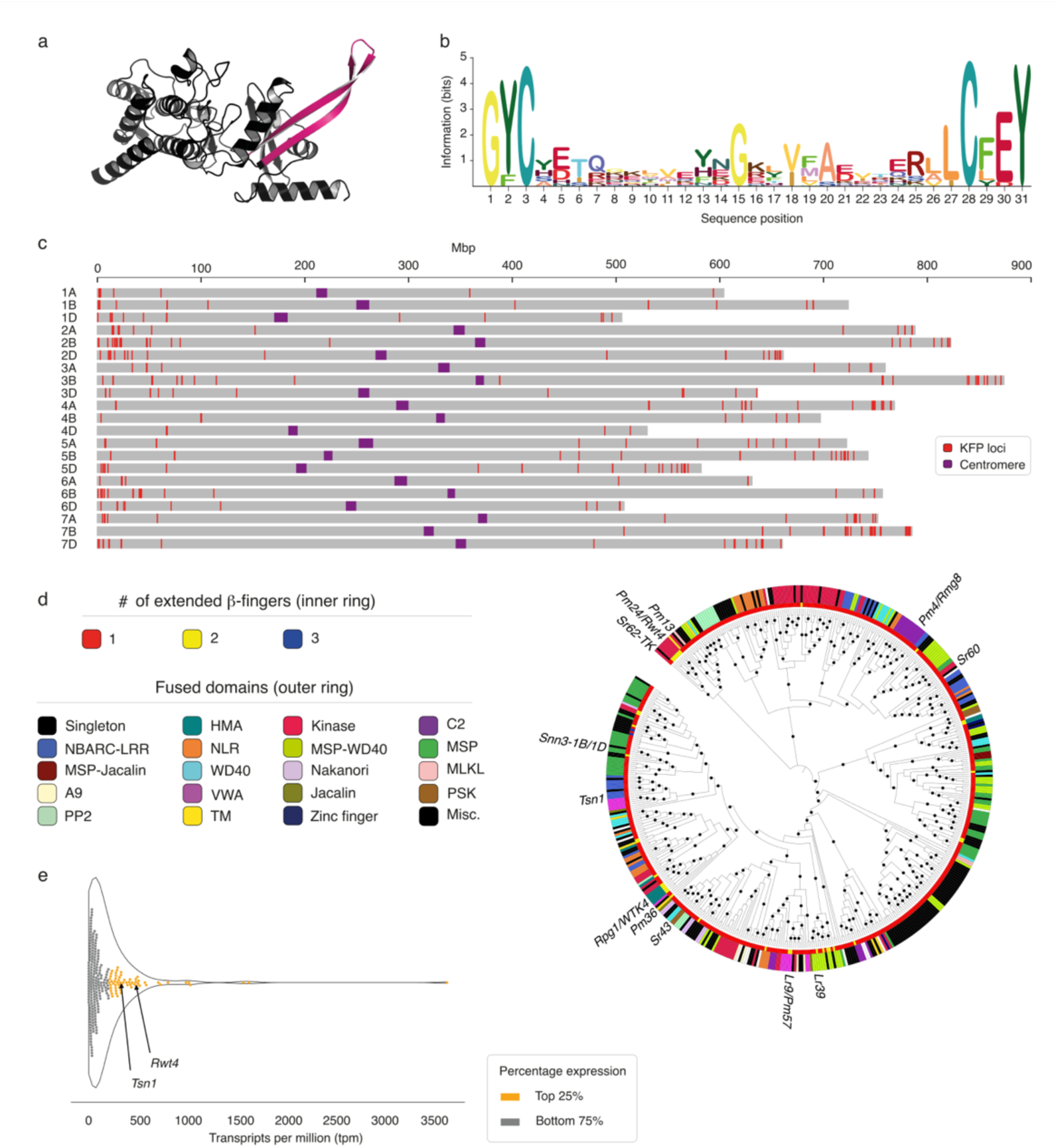
**a**, Predicted structure of the extended β-finger kinase domain of Sr62^TK^, showing the extended β-finger motif in magenta (Table S1). **b**, Sequence logo of the refined hidden Markov model (HMM) representing the extended β-finger in KFPs from wheat cv. Chinese Spring. **c**, Genomic distribution of KFP genes across the Chinese Spring genome. **d**, Phylogenetic tree based on the extended β-finger domain(s) (primary module) of all KFPs encoded by the Chinese Spring genome. Black circles indicate bootstrap value ≥80. KFPs derived from previously cloned genes were included and named. Sequences are coloured according to their number of extended β-finger kinase domains (inner circle) or auxiliary module (outer circle), as indicated in the legend. Auxiliary domains occurring fewer than three times across the phylogenetic tree are labelled miscellaneous (Misc.) (Table S4, S5). **e**, KFP gene transcript abundance in *de novo* assembled transcriptomes of wheat cv. Cadenza. The two functionally validated KFPs Tsn1 and Rwt4 are highlighted. Transcript levels were estimated from self-aligned RNA-seq data as transcripts per million (TPM). The top 25% most expressed KFP genes are shown in orange.

Using the refined HMM of the extended β-finger motif, we annotated the Chinese Spring proteome and identified 476 unique loci across the genome that encode a kinase domain with at least one such motif. These KFP loci were distributed across all chromosomes, with an enrichment in the gene-rich telomeric regions, similar to the published distribution of genes encoding nucleotide-binding leucine-rich repeat (NLR) resistance proteins (Fig. 1c, Table S4)^6^.

The number of KFP genes per chromosome ranged from 6 on chromosome 4D to 53 on chromosome 7B (Table S5).

To investigate the evolutionary relationships among KFPs, we extracted the extended β-finger kinase domains from the protein sequences encoded by each of the 476 unique KFP loci identified in Chinese Spring. We constructed a phylogenetic tree based on an alignment of the extended β-finger motif from these 476 KFPs and added the proteins encoded by previously cloned KFP genes as a reference (Fig. 1d, Table S1). We annotated the protein sequence of each KFP associated with an extended β-finger and identified 31 distinct auxiliary modules (i.e., fused domains) (Fig. 1d, Table S7). Only 10 of these 31 modules were previously reported in the proteins encoded by cloned KFP genes (Table S7). The most prevalent auxiliary modules included a non β-finger kinase domain (*n* = 59), a major sperm protein (MSP) domain (*n* = 53), an MSP domain combined with a WD40 repeat (MSP-WD40; *n* = 50), an NBARC-LRR domain (*n* = 29), and a coiled-coil NLR (CC-NLR; *n* = 25) (Table S7). The different clades of the KFP phylogenetic tree did not correlate with the subgenome origin of their encoding genes (Fig. S22).

When mapped onto the KFP phylogeny, the auxiliary modules formed distinct clades. In several cases, the same auxiliary module appeared in multiple, independently evolved clades, suggesting convergent domain acquisition (Fig. S23–24). Alternatively, perhaps the same acquisition events were followed by divergent evolution of their extended β-fingers. We also identified 150 KFPs with a kinase domain but lacking a detectable auxiliary domain (singletons), 55 of which were putatively annotated as cysteine-rich receptor-like kinases (crRLKs). These singletons were more frequent in several of the clades but were also dispersed throughout the phylogenetic tree (Fig. S25).

In addition, we identified 29 KFPs containing additional extended β-finger kinase domains, which clustered into clades associated with specific auxiliary modules (Fig. 1d) (Table S8). The most common auxiliary modules in these KFPs were kinase (*n* = 6), heavy metal– associated (HMA)-like (*n* = 6), and MSP-WD40 (*n* = 4) domains. In addition, we detected 11 tandem kinase domains with additional β-finger motifs but no associated auxiliary domain.

The functionally validated KFPs Sr62^TK^ and Rwt4 trigger immune responses via a shared CC-NLR, which is encoded by a locus located 20 kb distal and 150 kb proximal to the respective KFP genes. Therefore, we investigated whether other KFP genes in the Chinese Spring genome were also located close to CC-NLR genes: indeed, 28.2% of KFP genes were located within 200 kb of a CC-NLR gene (Fig. S26, Table S9).

Most NLR genes validated as resistance genes—across both dicots and monocots—are among the top 25% most highly expressed NLR genes within a plant genome’s NLR repertoire^7^. To assess whether those KFPs conferring disease resistance are encoded by highly expressed genes, we quantified the transcript levels of 12 KFP genes using publicly available transcriptome deep sequencing (RNA-seq) datasets (Table S10). These previously cloned KFP genes consistently ranked among the top 25% of expressed KFP genes across each dataset, with the exception of *Rpg1* (Fig. 1e, S27a–i). Moreover, the *Sr62^TK^*and *Sr62^NLR^* pair were expressed at similar levels and were both in the top 25% of the most highly expressed KFP and NLR genes, respectively (Fig. S27h and S28).

To investigate the evolutionary origin of the extended β-finger in plants, we annotated the motif in available reference genomes for monocot species. The extended β-finger motif was present exclusively in members of the Poaceae family, except for the early-diverging grass *Streptochaeta angustifolia* (Fig. 2, Table S11). This pattern suggests that the motif emerged at least 98 million years ago, following the divergence of *S. angustifolia* and *Pharus latifolius*^8^. The apparent absence of the motif within the Poaceae species *Avena byzantina* and *Neostapfia colusana* may reflect intraspecific variation or species-specific loss.

**Fig 2.**
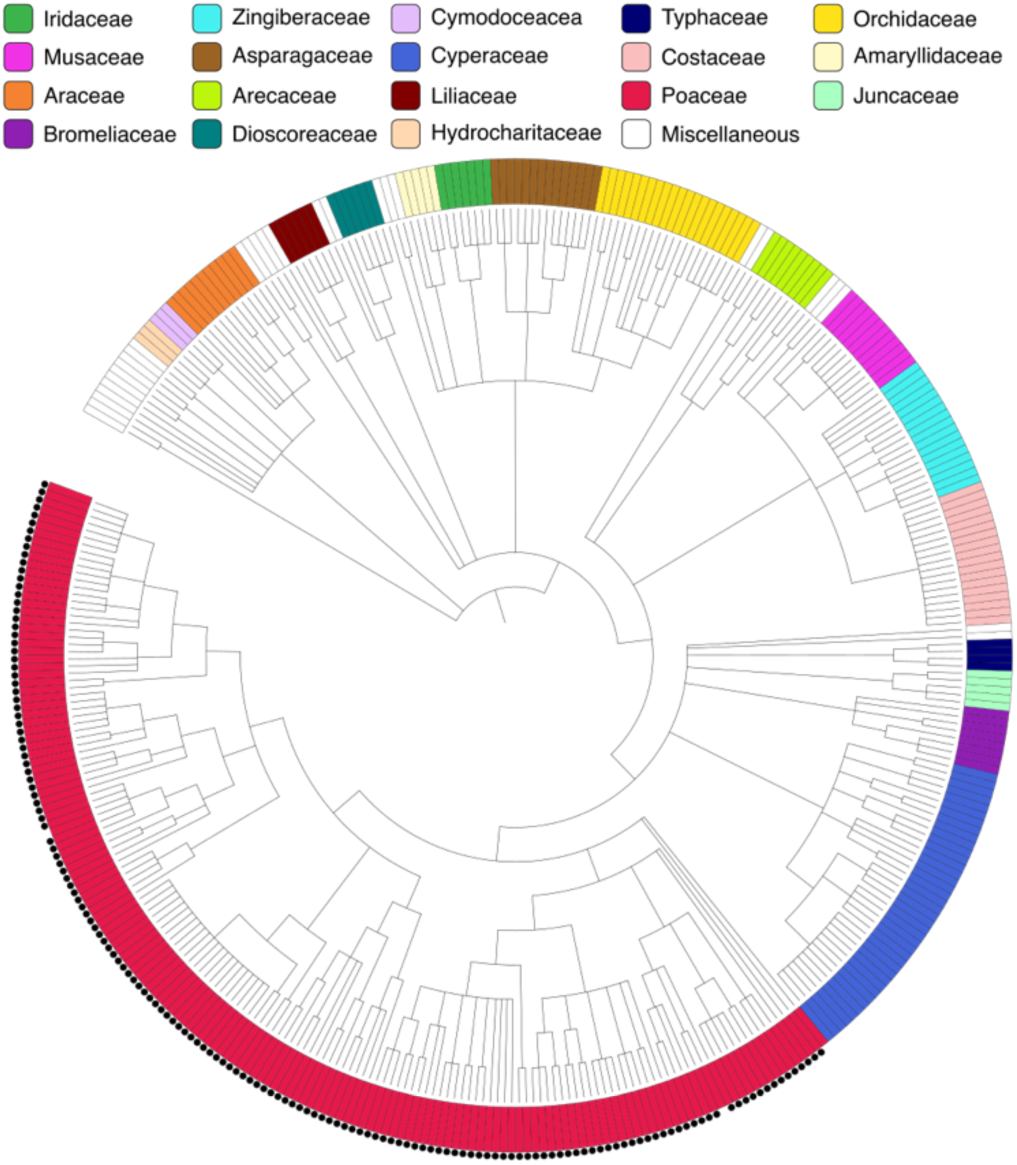
Extended β-finger KFPs are unique to the Poaceae family. Presence or absence of the extended β-finger motif in publicly available genomes of monocotyledonous species. A black circle indicates a species in which the motif was detected. Members of each family are highlighted according to the legend. Families with fewer than three members in the tree are labelled as miscellaneous. Species names can be found in Supplementary Table 11.

Through a computational analysis of cloned KFP genes, we demonstrated that the extended β-finger is a highly conserved motif that can be used for the *de novo* annotation of immune receptors as KFPs. We believe this knowledge will be useful for the molecular cloning of as yet unknown resistance genes from Poaceae genomes, as KFP genes comprise ∼20% of currently cloned resistance genes in wheat ^9^. Our analysis of the wheat cv. Chinese Spring genome identified 476 unique KFP loci; however, as most functionally validated KFPs are typically encoded by highly expressed genes, not all identified KFPs will likely confer disease resistance^6,7^.

Across the phylogeny of KFPs from Chinese Spring, distinct clades share the same auxiliary modules. For example, KFPs with an MSP or a kinase auxiliary module form multiple distinct clades. The low sequence similarity between the kinase domains of these clades suggests that KFPs in the different clades likely arose from distinct independent fusions, whereas those within a single clade may share the same ancestral fusion event. Given their abundance and presence across many clades in the tree, crRLKs are likely the source of the kinase domains involved in these fusion events, suggesting that KFPs are the result of multiple independent fusion events. The identified singleton kinase domains may represent the true pathogenicity targets of the effectors recognized by the KFPs in these clades^10^. Alternatively, the auxiliary modules fused to the kinase domains may represent integrated decoys for homologous effector targets, as is the case for NLRs with integrated domains ^11^.

Although KFP genes have been cloned exclusively from barley, wheat, and wild wheat relatives, our phylogenetic analysis of monocot proteomes highlights a potentially untapped wealth of KFP resistance genes available in other members of the Poaceae such as rice (*Oryza sativa*), maize (*Zea mays*), and their wild relatives. Our analysis also highlights the specificity of the extended β-finger motif to the Poaceae. However, since crRLKs are ubiquitous across plant species, it is possible that KFPs lacking the extended β-finger motif, as exemplified by the tandem kinase Yr15, occur outside the Poaceae.

## Methods

### Identification of the extended β-finger via HMM

AlphaFold 3 (ref.^12^) was used to predict the structures of cloned KFPs (Table 1). The quality of each prediction was assessed using AlphaFold Analyser (v3.1.0). The resulting structures were visualized in PyMol and manually curated to identify and extract the sequence of the extended β-finger motif. A multiple sequence alignment (MSA) of unique extended β-finger sequences was generated using MUSCLE v5.1 (ref.^13^). The MSA was used to create a seed hidden Markov model (HMM) using *hmmbuild* with default parameters ^14^.

The seed HMM was then iteratively refined against the reference proteome of wheat cultivar Chinese Spring^15^. At each iteration step, *hmmsearch*, with the parameters *–incdome 1e-10* and *-A proteome.aln*, was used to identify and align protein sequences containing a β-finger^14^. The number of unique sequences was counted and, using the alignment, a new HMM was constructed using *hmmbuild*. The newly created HMM was then used for the next iteration step^14^. This process was continued until three successive iteration steps identified the same number of proteins in the proteome. The number of sequences annotated in each refinement step was plotted using plot_refinement.py (https://github.com/Orpowell/wheat_kfps). The matrix for the refined HMM was extracted using Skylign, and the sequence logo of the HMM was generated with plot_sequence_logo.py (https://github.com/Orpowell/wheat_kfps) ^16^.

### Chinese Spring annotation and phylogenetics

A proteome of the telomere-to-telomere (T2T) assembly of the Chinese Spring genome was generated by selecting the longest protein sequence for each annotated locus using get_locus_representatives.py (https://github.com/Orpowell/wheat_kfps) ^15^. The resulting non-redundant proteome was independently annotated for extended β-fingers motifs and Pkinase domains using *hmmsearch,* with the parameter --domE 1e-4, and the extended β-finger HMM (see above) and the protein kinase (Pkinase) HMM from PFAM, respectively ^14,17^. A protein was classified as a KFP if it contained at least one extended β-finger overlapping with the Pkinase domain. This analysis was performed using “annotate_beta_finger_and_kinase.sh” (https://github.com/Orpowell/wheat_kfps).

The primary module, defined here as the extended β-finger kinase domain(s), was extracted from the KFPs in the Chinese Spring proteome and from previously cloned KFPs using extract_kfp_sequences.py (https://github.com/Orpowell/wheat_kfps) and aligned using MUSCLE v5.1 with default parameters ^13^. The resulting multiple sequence alignment was used to reconstruct a phylogenetic tree using FastTree (v2.1.11) with default parameters ^18^. The multiple sequence alignment was also used to generate a percentage identity matrix (PIM) of KFPs using Clustal omega^19^ with the parameters: “--percent-id --distmat-out=PIM.txt –full -- force”.

The phylogenetic tree was visualized using iTOL v7 ^20^. An iTOL annotation for the number of β-fingers in each KFP was generated using count_beta_fingers.py (https://github.com/Orpowell/wheat_kfps). Protein sequences of Chinese Spring KFPs and previously cloned KFPs (Table S1) were annotated using InterProScan, with TMHMM enabled ^21,22^. The auxiliary modules in each KFP were identified and used to create an iTOL annotation using annotate_KFP_auxiliary_domain.py (https://github.com/Orpowell/wheat_kfps). This annotation file was filtered to create specific annotations for KFPs with MSP and kinase auxiliary modules. PIMs for the extended β-finger kinase domain(s) of KFPs with MSP and kinase auxiliary modules were generated using “analyse_pim.py” (https://github.com/Orpowell/wheat_kfps).

The coordinates of KFP genes were extracted from the Chinese Spring reference GFF file and visualized using plot_KFP_physical_positions.py (https://github.com/Orpowell/wheat_kfps). The subgenome location of each KFP gene was extracted and annotated using annotate_subgenome.py (https://github.com/Orpowell/wheat_kfps).

### KFP–NLR proximity in Chinese Spring

NLR loci in the T2T Chinese Spring genome were annotated using NLR-Annotator with default parameters ^23^. The nearest CC-NBARC-LRR locus to each KFP locus (identified above) was determined using calculate_nlr_proximity.py (https://github.com/Orpowell/wheat_kfps).

### HMM-Annotator

HMM-Annotator is a Python tool for the annotation of genome and transcriptome sequences using protein HMMs written specifically for this study. The tool works via the following steps. First, the input DNA sequence is dissected into overlapping fragments. Each fragment is then fully translated into its six possible protein sequences (three forward and three reverse open reading frames) that are annotated for HMM using PyHMMer. Protein sequence annotations are then converted back to their original DNA sequence coordinates. The output can be saved as a TSV or a BED file. HMM-Annotator was written in Python (v3.10.14). Source code and documentation can be found on GitHub (https://github.com/Orpowell/HMM-Annotator).

### Expression analysis of cloned KFP genes

Publicly available RNA-seq data for cloned KFP genes were downloaded from NCBI and NGDC (Table S10). The raw reads from each dataset were trimmed to remove adapters and low-quality sequences using Trimmomatic (v.0.39) with the parameters: LEADING:5,TRAILING:5, SLIDINGWINDOW:4:20, MINLEN:50 (ref.^24^). Trimmed reads were *de novo* assembled using Trinity (v2.15.2) using default parameters^25^. Transcript abundance was estimated using Salmon (v1.10.3), with bias correction enabled (--gcBias, -- seqBias), and aggregated into gene-level transcript per million (TPM) counts using the R package tximport (v1.36.0) ^26,27^. The transcripts of genes encoding proteins containing a β-finger were identified using HMM-Annotator (as described above), and their corresponding TPM values were calculated. Transcripts corresponding to cloned KFP genes were identified using BLASTn ^28^. KFP gene expression data were visualized using TEA-tools visualize command (https://github.com/Orpowell/tea-tools). Analysis of *Sr62^NLR^*was performed as described above for KFP genes, except that NLR-Annotator^23^ was used to identify NLR transcripts. The GenBank sequence PP537390.1 was used for the annotation of *Sr62^NLR^*transcripts.

### KFP annotation in monocot genomes

Publicly available reference genomes of monocot species were downloaded from NCBI using the NCBI datasets command: “datasets download genome taxon 4447 –dehydrated –reference –include genome”. The genome of *Streptochaeta angustifolia* was downloaded using the NCBI datasets command: “download genome accession GCA_020804685.1 –include genome”. The name of each species was extracted from the “assembly_report.jsonl” included in the data downloads using get_species_name.py (https://github.com/Orpowell/wheat_kfps). A phylogenetic tree of species was downloaded from NCBI using ete3 and visualized using iTOL v7 ^20,29^.

Each genome was annotated for extended β-finger motifs using HMM-Annotator and the extended β-finger motif HMM (see above). The output BED file for each genome was filtered for annotations with a minimum length of 90 nucleotides and score (E-value) less than 1E−10. Presence or absence of the motif was determined from the filtered BED file and used to generate an iTOL annotation for the phylogenetic tree using generate_itol_annotations.py (https://github.com/Orpowell/wheat_kfps).

Taxa information for each monocot species was downloaded using the NCBI datasets commands: “datasets summary taxonomy taxon 4447 --as-json-lines --children | dataformat tsv taxonomy --template tax-summary”. The resulting TSV file was used to generate an iTOL annotation of monocot families present in the phylogenetic tree using generate_family_annotation.py (https://github.com/Orpowell/wheat_kfps).

## Acknowledgements

This research used the Ibex high-performance computing cluster managed by the Supercomputing Laboratory at King Abdullah University of Science and Technology (KAUST), Saudi Arabia. We are grateful to Burkhard Steuernagel for helpful discussions during the development of HMM-Annotator and Tobin Florio (https://flozbox-science.com) for figure artwork.

## Author Contributions

O.R.P. developed the β-finger HMM and HMM-Annotator. O.R.P. performed all bioinformatic analyses related to Chinese Spring and the evolutionary origin of the extended β-finger motif. O.R.P. developed HMM-Annotator and TEA-tools. D.G. performed the expression analysis of cloned KFP genes. B.B.H.W. and O.R.P. conceived the study. O.R.P. drafted the manuscript with contributions from B.B.H.W. and D.G. All coauthors read and approved the final manuscript.

## Data and Materials Availability

Source code for HMM-Annotator, Tea-tools, and all analyses are available on GitHub: (https://github.com/Orpowell/HMM-Annotator), (https://github.com/Orpowell/tea-tools), (https://github.com/Orpowell/wheat_kfps). AlphaFold3 models used to identify the extended β-finger motif are archived on Zenodo (https://doi.org/10.5281/zenodo.16732312). The HMM for the extended β-finger motif is archived on Zenodo (https://doi.org/10.5281/zenodo.15874664).

## Supplementary Figures

**Supplementary Fig. 1.**
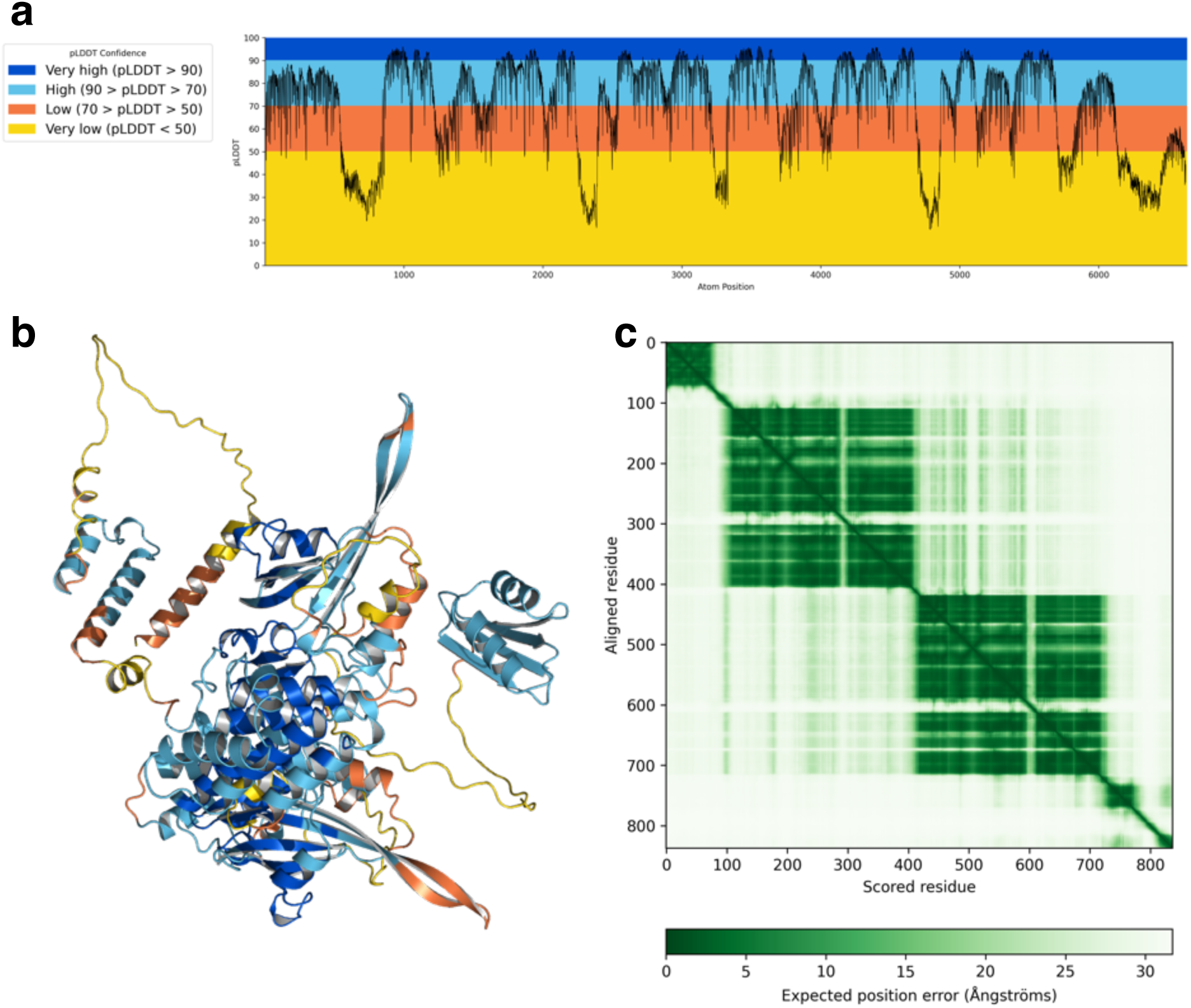
AlphaFold3 Prediction of Rpg1. pLDDT plot (**a**), pLDDT coloured structure (**b**), and predicted aligned error plot (**c**) for the highest confidence prediction of Rpg1 (Average pLDDT = 70.18, pTM = 0.43).

**Supplementary Fig. 2.**
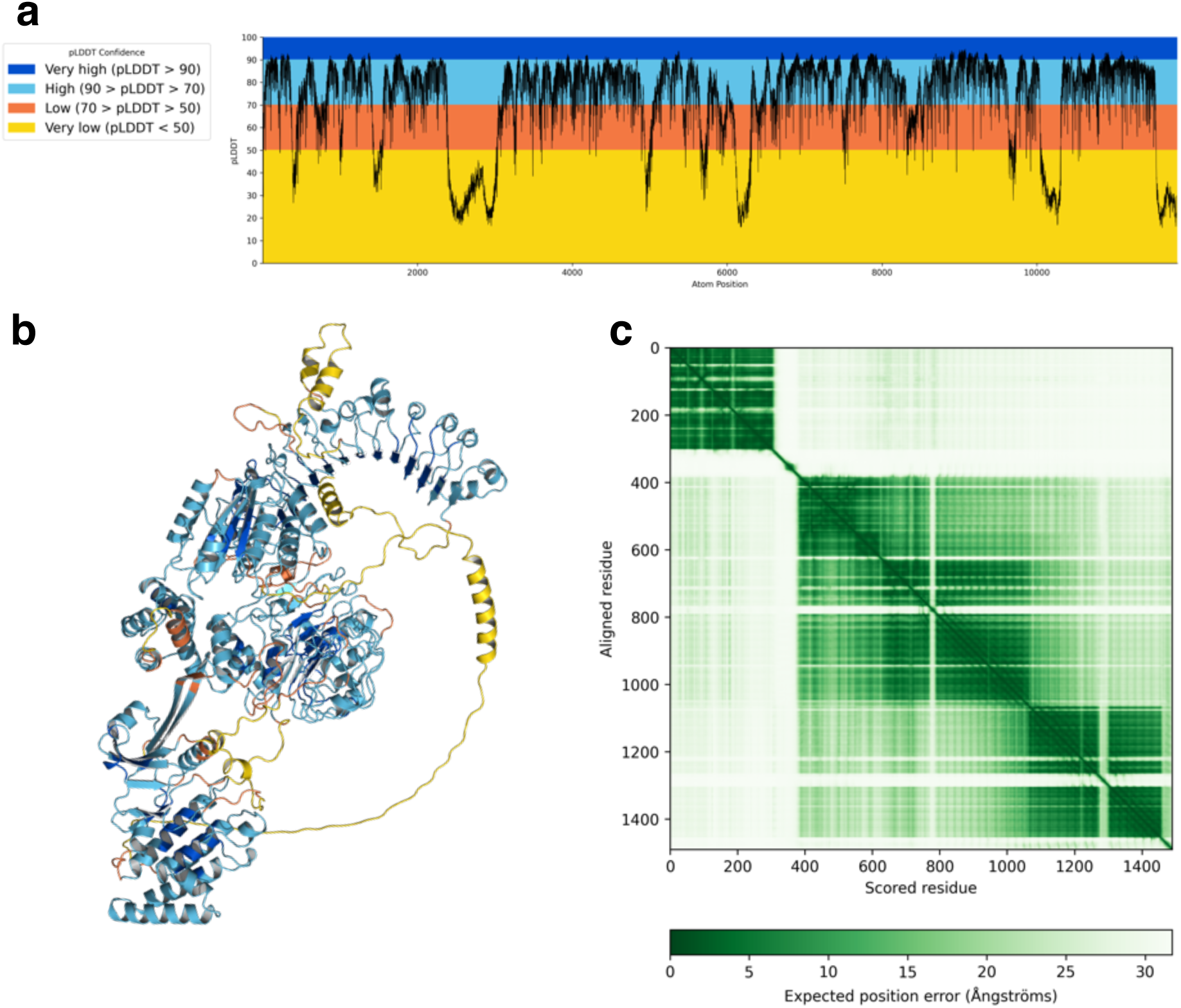
AlphaFold3 Prediction of Tsn1. pLDDT plot (**a**), pLDDT coloured structure (**b**), and predicted aligned error plot (**c**) for the highest confidence prediction of Tsn1 (Average pLDDT = 72.31, pTM = 0.54).

**Supplementary Fig. 3.**
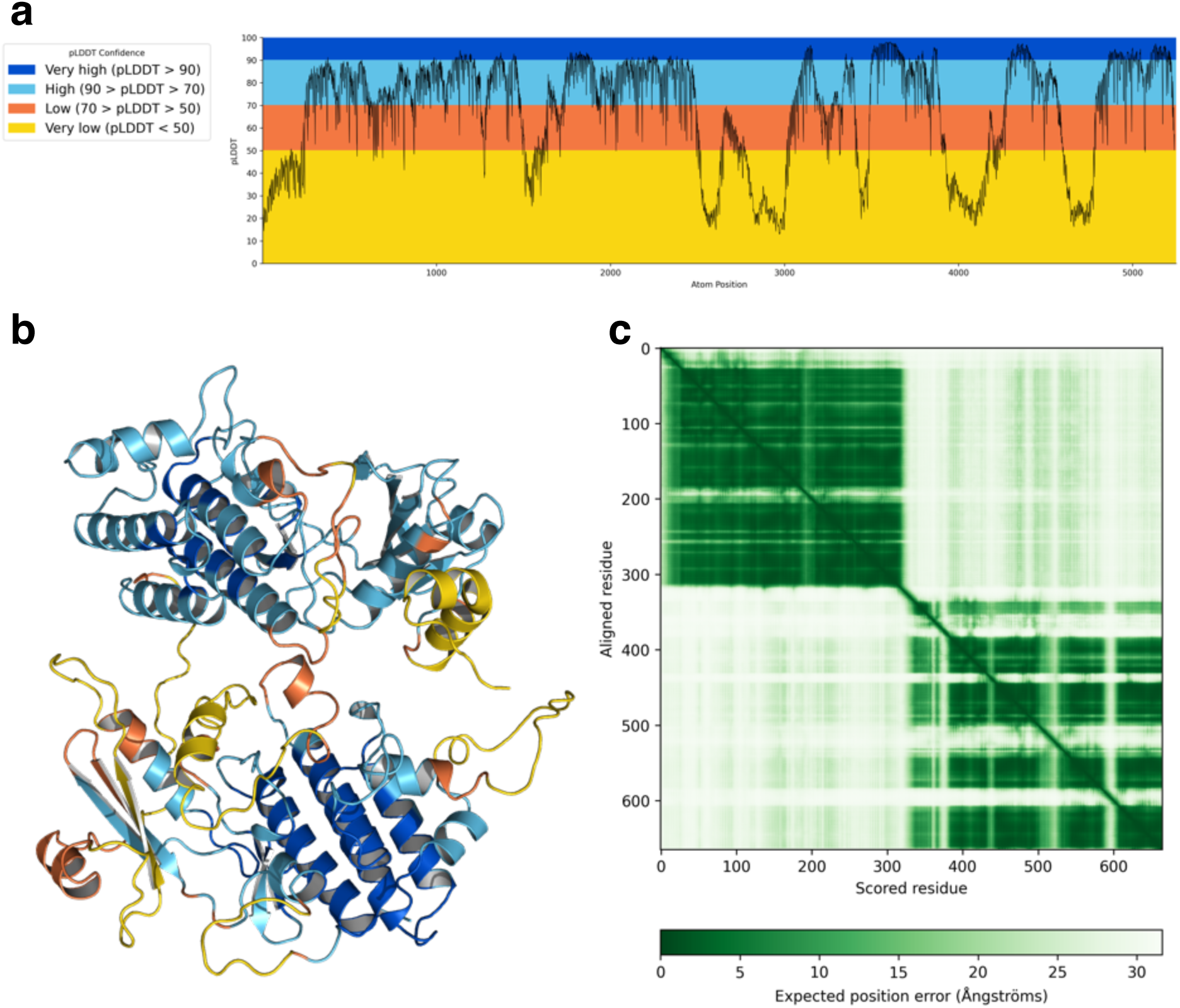
AlphaFold3 Prediction of Yr15. pLDDT plot (**a**), pLDDT coloured structure (**b**), and predicted aligned error plot (**c**) for the highest confidence prediction of Yr15 (Average pLDDT = 67.79, pTM = 0.51).

**Supplementary Fig. 4.**
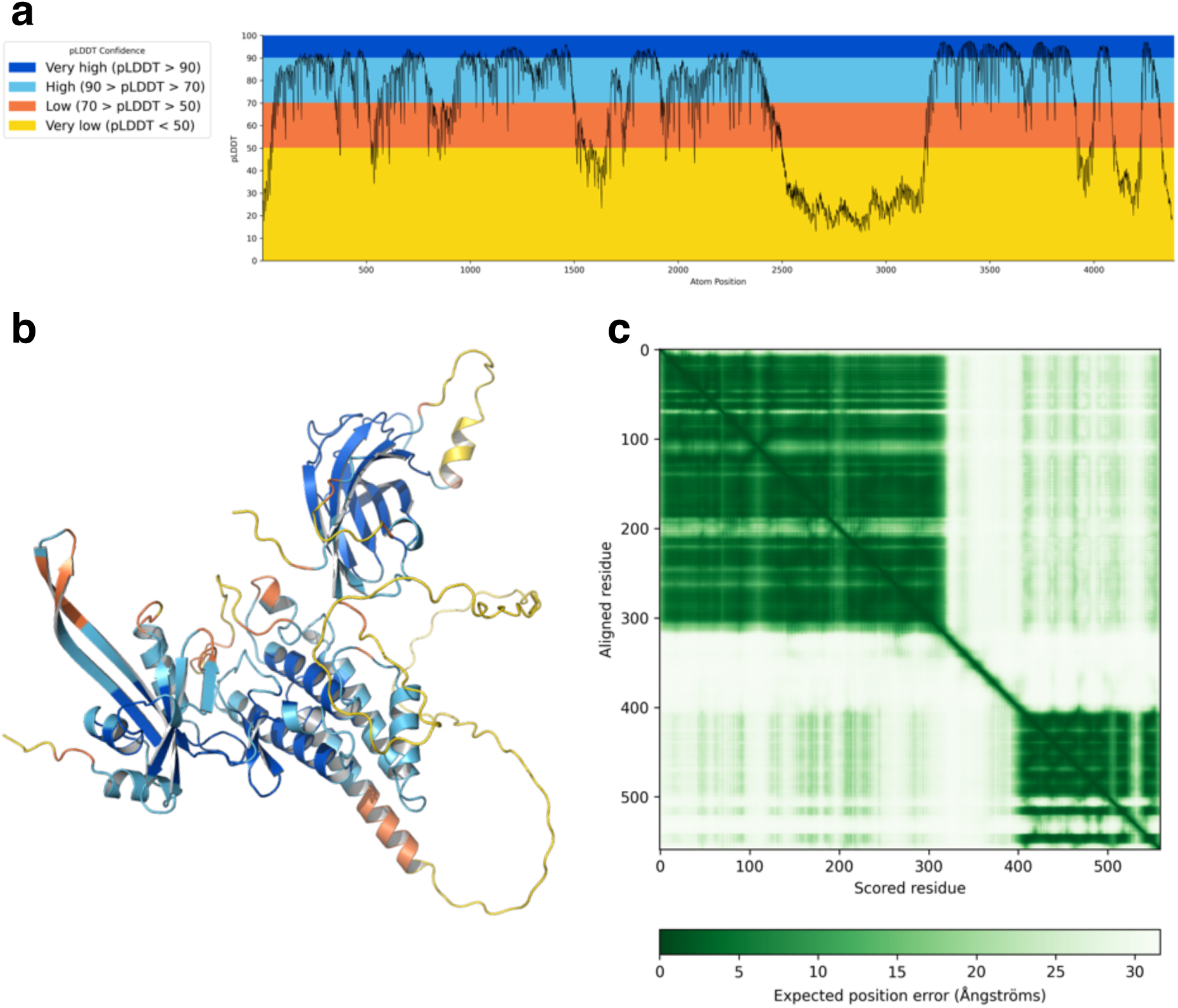
AlphaFold3 Prediction of Pm4d_v1. pLDDT plot (**a**), pLDDT coloured structure (**b**), and predicted aligned error plot (**c**) for the highest confidence prediction of Pm4d_v1 (Average pLDDT = 68.81, pTM = 0.58).

**Supplementary Fig. 5.**
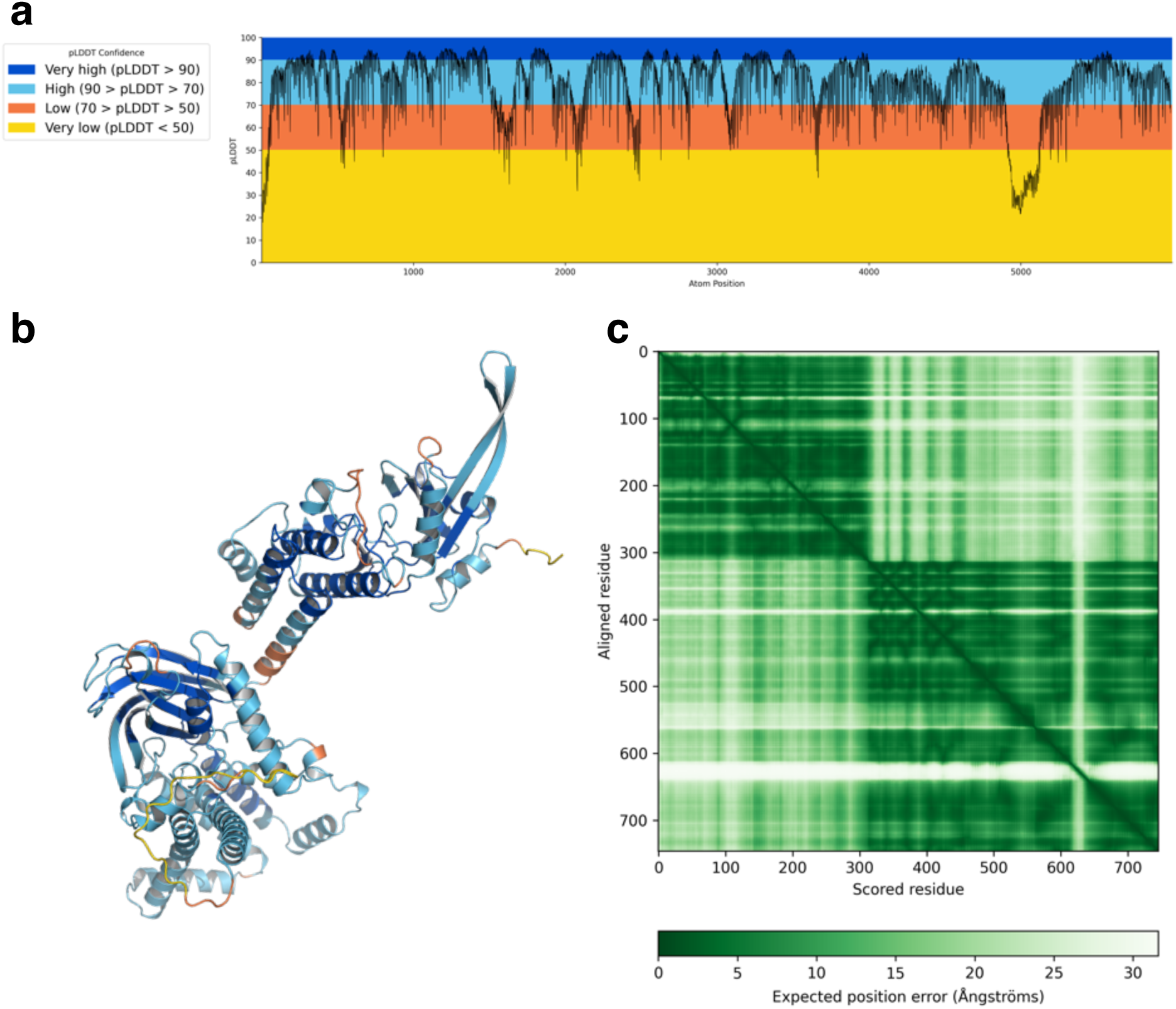
AlphaFold3 Prediction of Pm4d_v2. pLDDT plot (**a**), pLDDT coloured structure (**b**), and predicted aligned error plot (**c**) for the highest confidence prediction of Pm4d_v2 (Average pLDDT = 78.47, pTM = 0.67).

**Supplementary Fig. 6.**
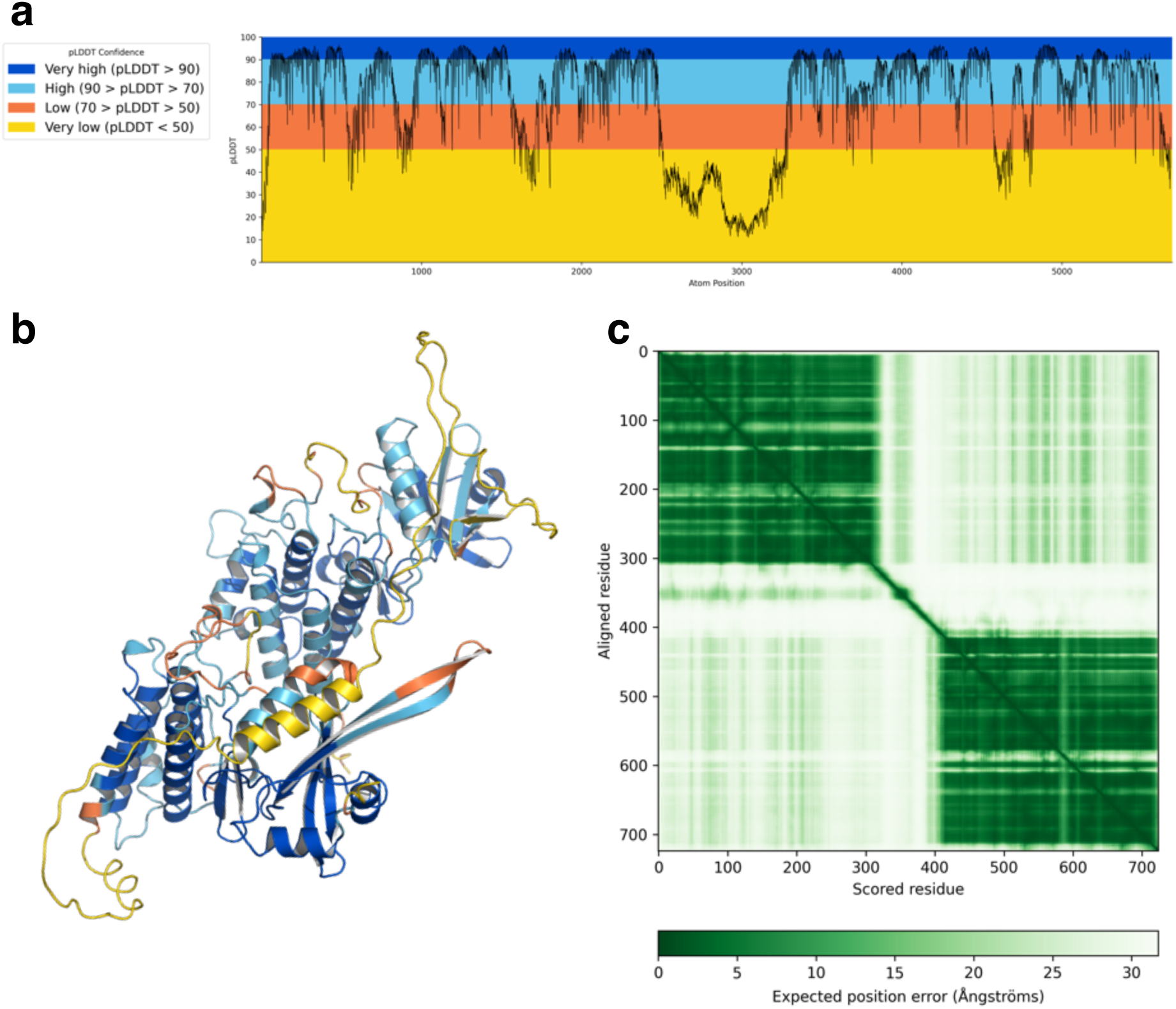
AlphaFold3 Prediction of Sr60. pLDDT plot (**a**), pLDDT coloured structure (**b**), and predicted aligned error plot (**c**) for the highest confidence prediction of Sr60 (Average pLDDT = 72.92, pTM = 0.51).

**Supplementary Fig. 7.**
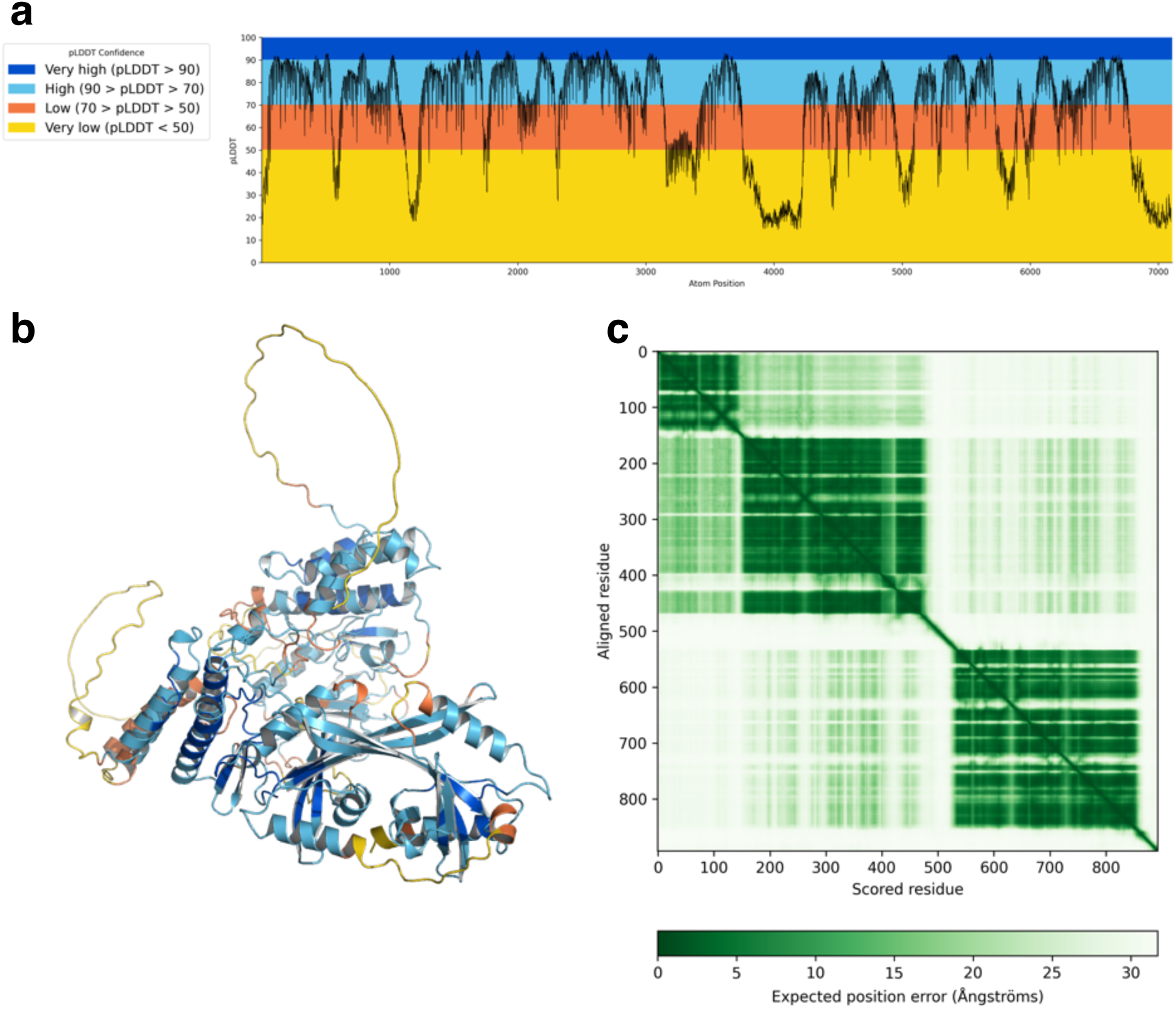
AlphaFold3 Prediction of Pm24. pLDDT plot (**a**), pLDDT coloured structure (**b**), and predicted aligned error plot (**c**) for the highest confidence prediction of Pm24 (Average pLDDT = 67.95, pTM = 0.49).

**Supplementary Fig. 8.**
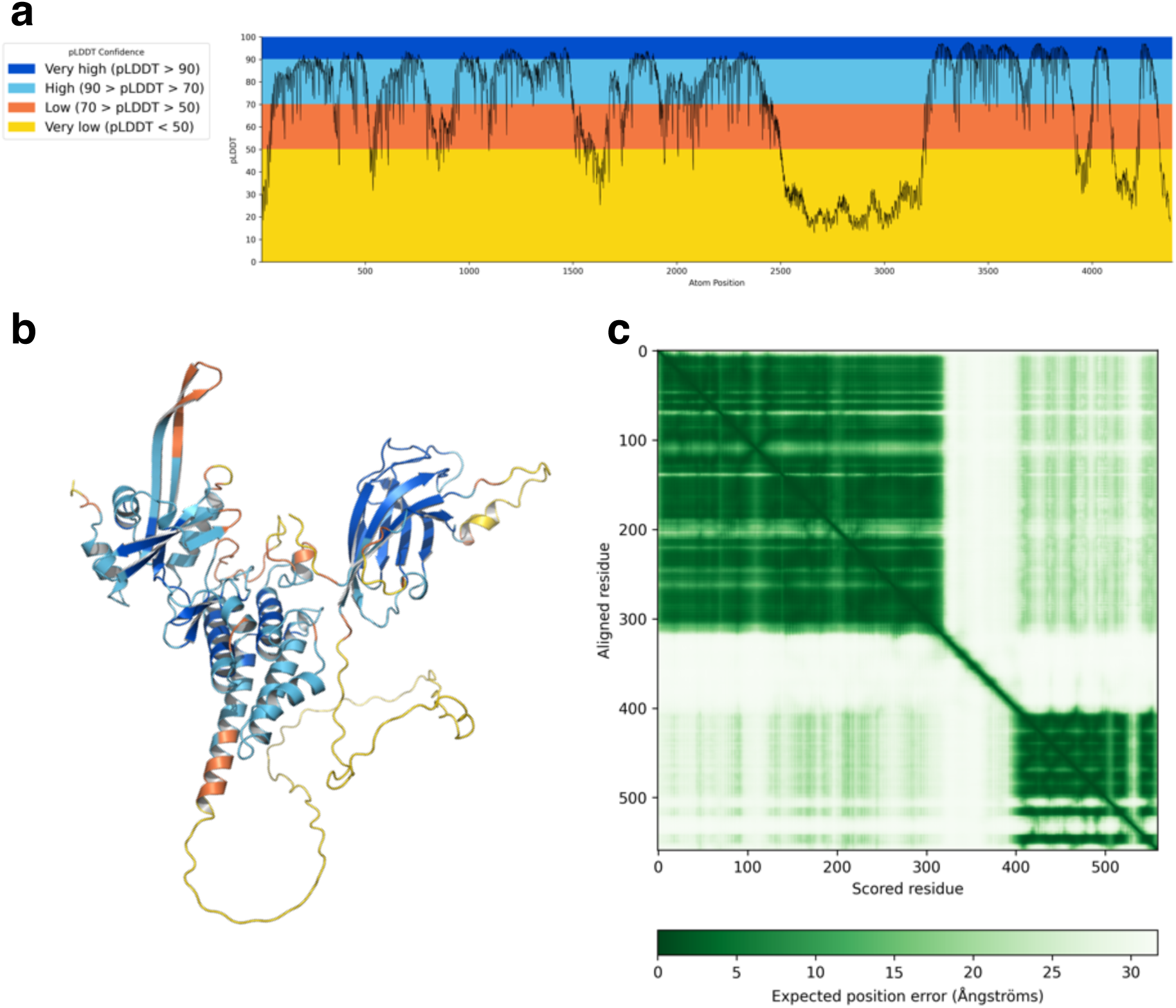
AlphaFold3 Prediction of Pm4b_v1. pLDDT plot (**a**), pLDDT coloured structure (**b**), and predicted aligned error plot (**c**) for the highest confidence prediction of Pmb4b_v1 (Average pLDDT = 68.26, pTM = 0.59).

**Supplementary Fig. 9.**
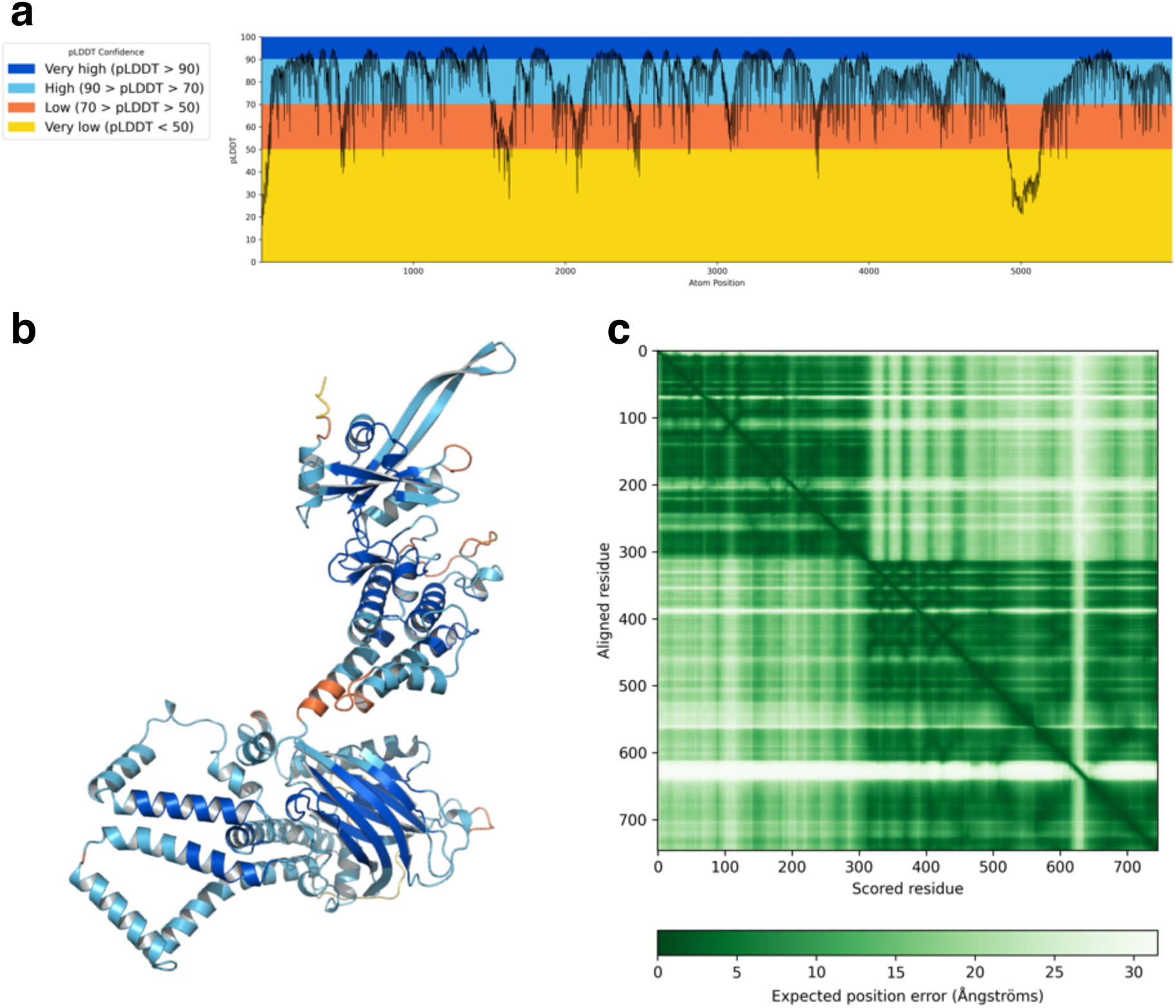
AlphaFold3 Prediction of Pm4b_v2. pLDDT plot (**a**), pLDDT coloured structure (**b**), and predicted aligned error plot (**c**) for the highest confidence prediction of Pm4b_v2 (Average pLDDT = 78.19, pTM = 0.68).

**Supplementary Fig. 10.**
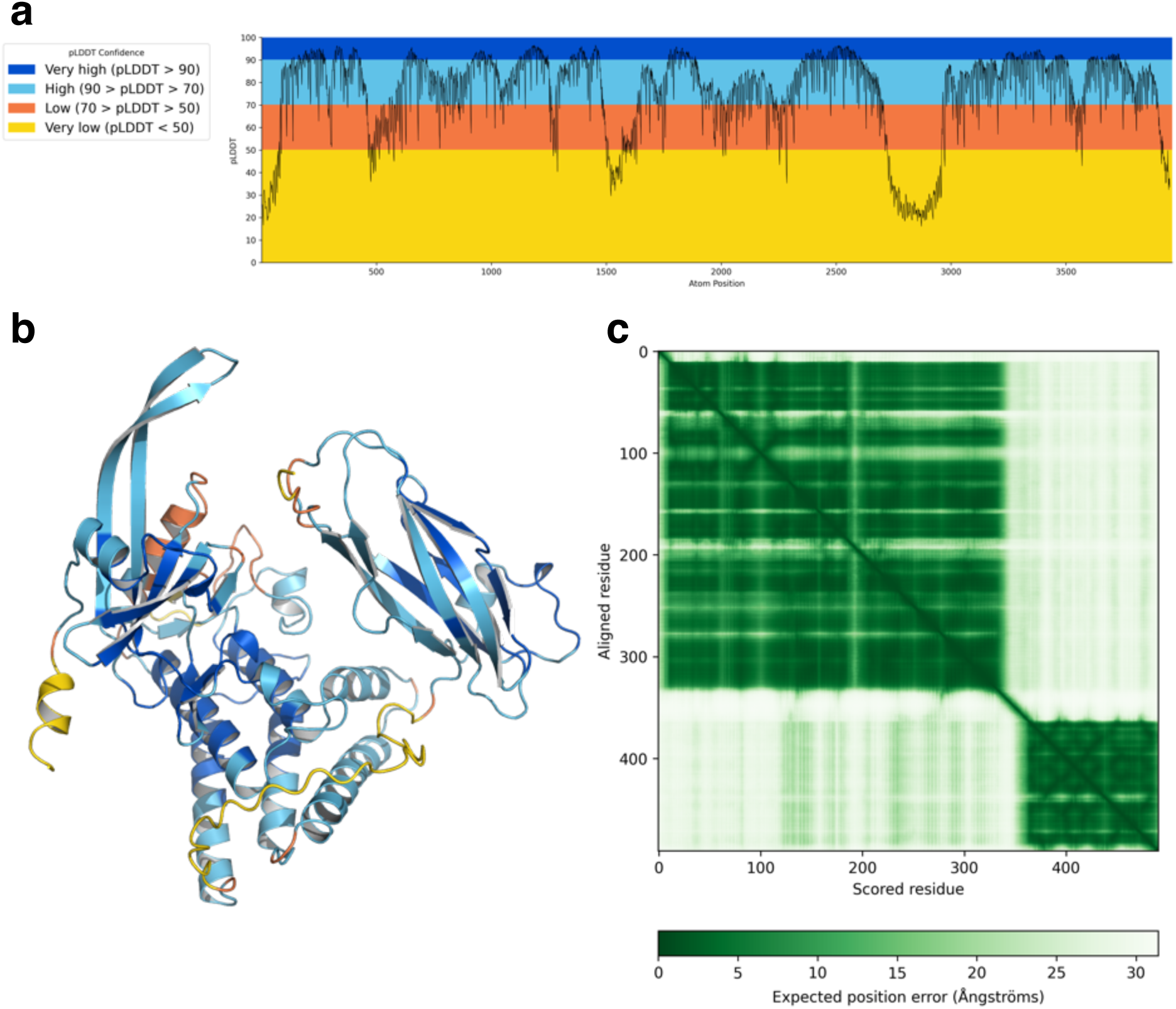
AlphaFold3 Prediction of Snn3-1D. pLDDT plot (**a**), pLDDT coloured structure (**b**), and predicted aligned error plot (**c**) for the highest confidence prediction of Snn3-1D (Average pLDDT = 75.24, pTM = 0.65).

**Supplementary Fig. 11.**
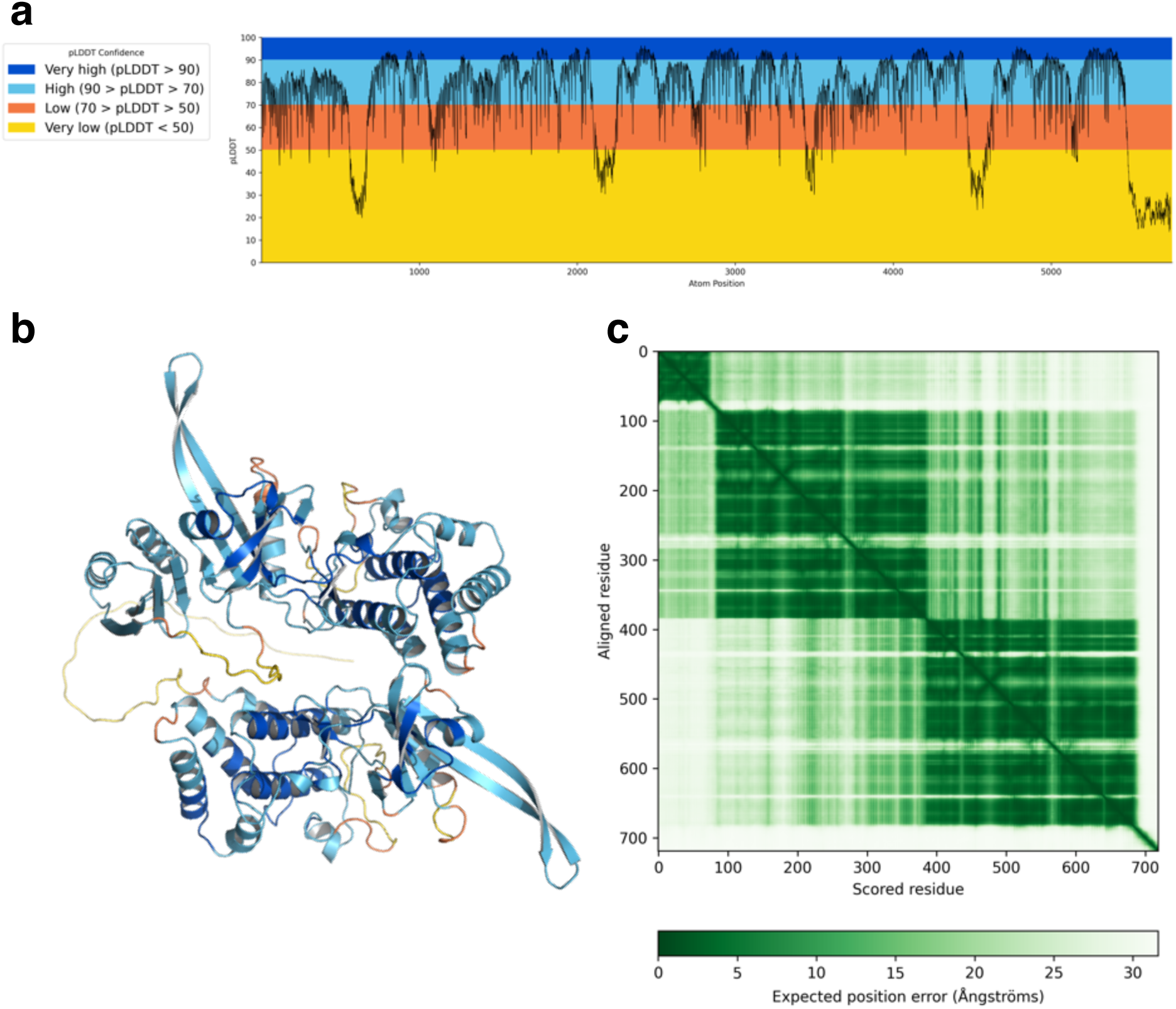
AlphaFold3 Prediction of WTK4. pLDDT plot (**a**), pLDDT coloured structure (**b**), and predicted aligned error plot (**c**) for the highest confidence prediction of WTK4 (Average pLDDT = 74.74, pTM = 0.58).

**Supplementary Fig. 12.**
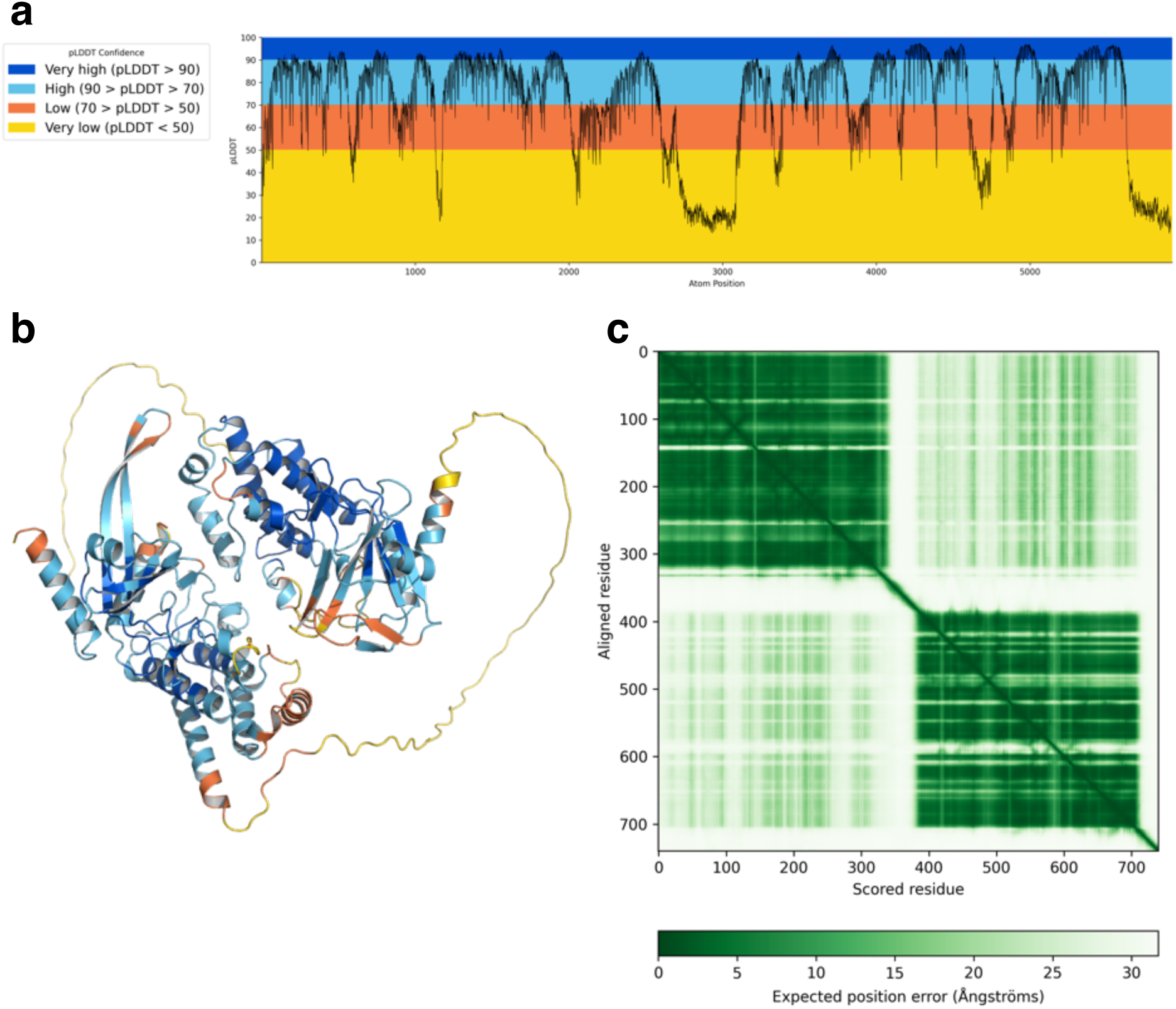
AlphaFold3 Prediction of Sr62. pLDDT plot (**a**), pLDDT coloured structure (**b**), and predicted aligned error plot (**c**) for the highest confidence prediction of Sr62 (Average pLDDT = 71.31, pTM = 0.54).

**Supplementary Fig. 13.**
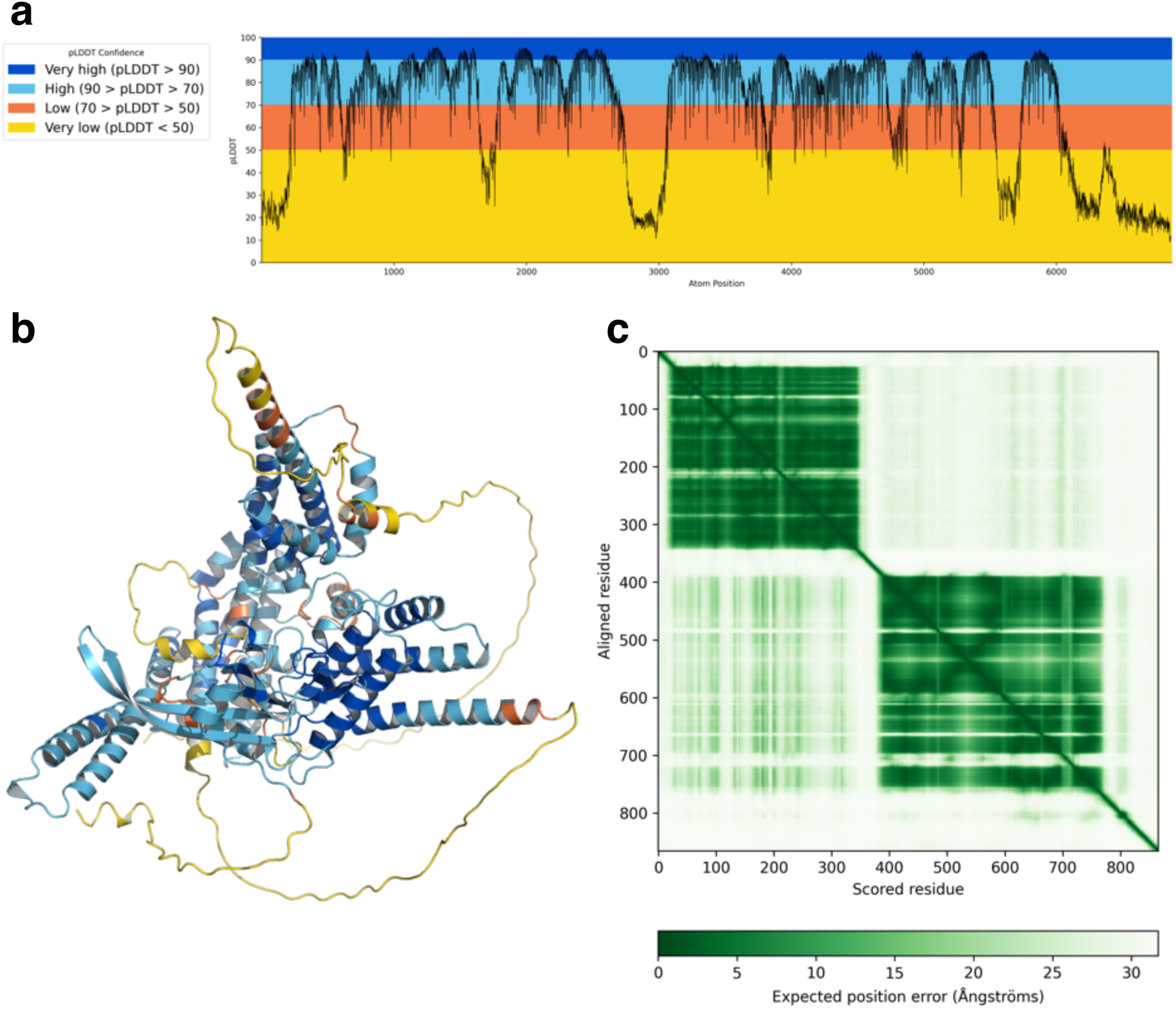
AlphaFold3 Predictions of Sr43. pLDDT plot (**a**), pLDDT coloured structure (**b**), and predicted aligned error plot (**c**) for the highest confidence prediction of YYY (Average pLDDT = 67.80, pTM = 0.48).

**Supplementary Fig. 14.**
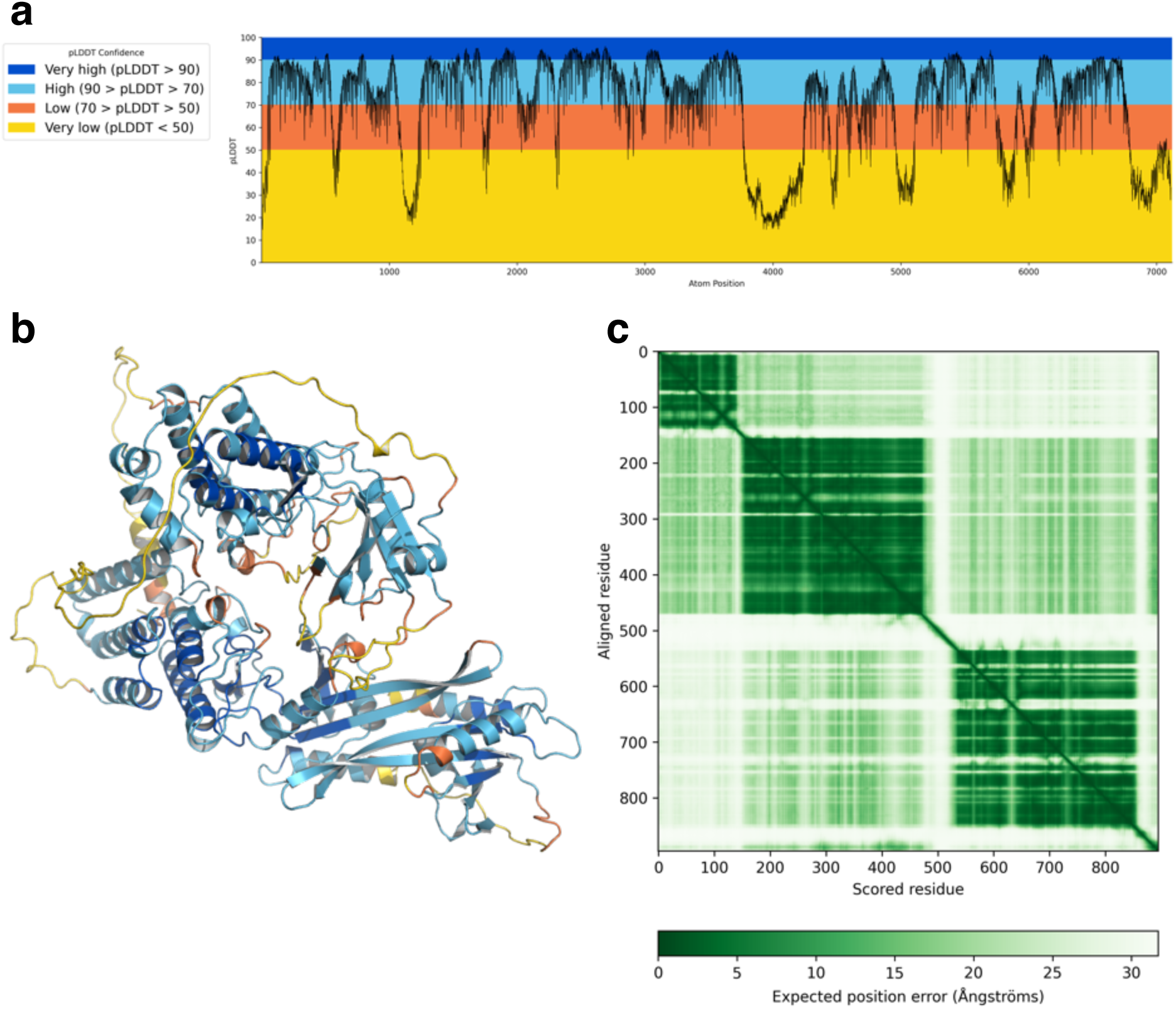
AlphaFold3 Prediction of Rwt4. pLDDT plot (**a**), pLDDT coloured structure (**b**), and predicted aligned error plot (**c**) for the highest confidence prediction of Rwt4 (Average pLDDT = 70.25, pTM = 0.55).

**Supplementary Fig. 15.**
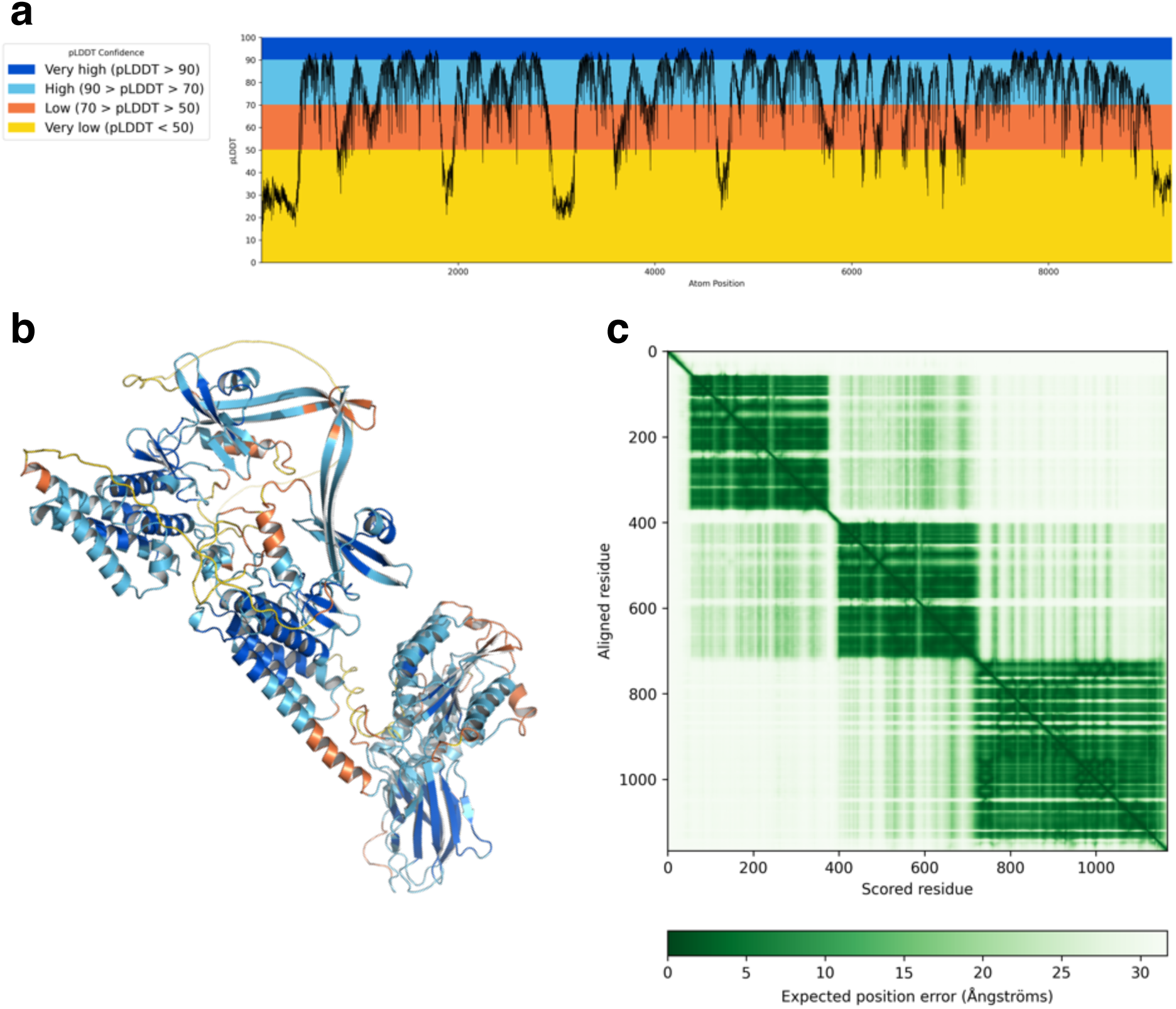
AlphaFold3 Prediction of Lr9. pLDDT plot (**a**), pLDDT coloured structure (**b**), and predicted aligned error plot (**c**) for the highest confidence prediction of Lr9 (Average pLDDT = 72.54, pTM = 0.45).

**Supplementary Fig. 16.**
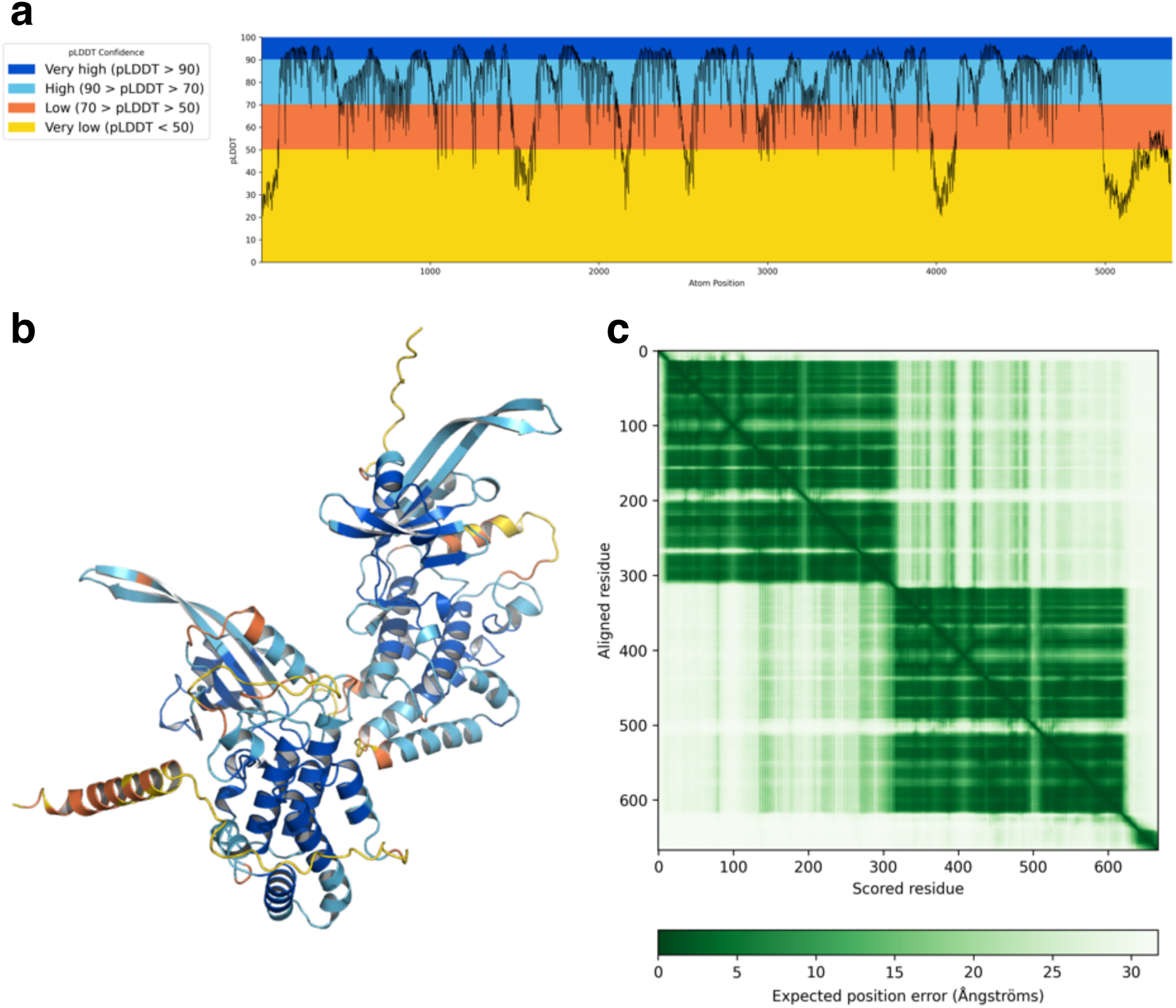
AlphaFold3 Prediction of Pm36. pLDDT plot (**a**), pLDDT coloured structure (**b**), and predicted aligned error plot (**c**) for the highest confidence prediction of Pm36 (Average pLDDT = 0.52, pTM = 75.92).

**Supplementary Fig. 17.**
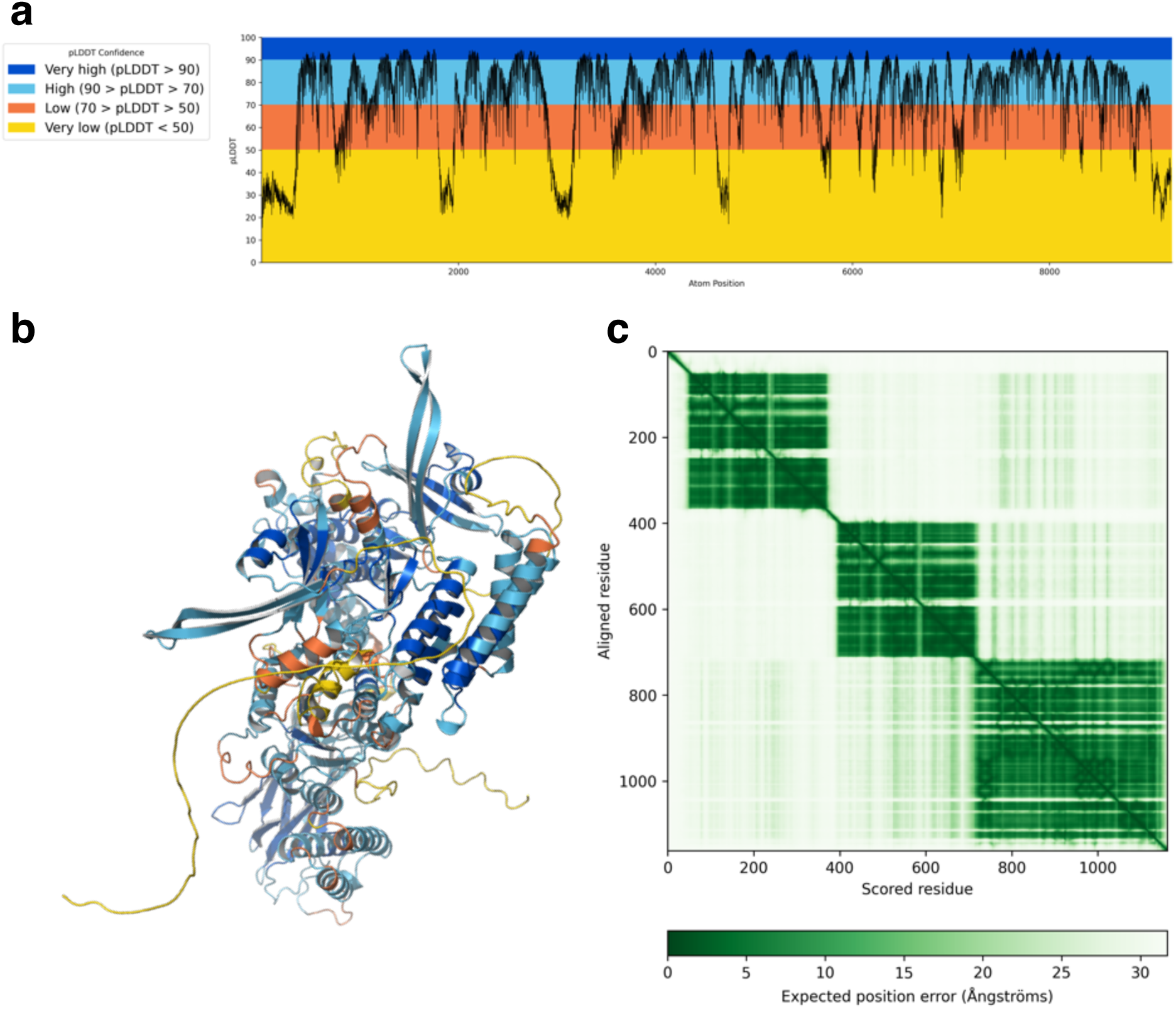
AlphaFold3 Prediction of Pm57. pLDDT plot (**a**), pLDDT coloured structure (**b**), and predicted aligned error plot (**c**) for the highest confidence prediction of Pm57 (Average pLDDT = 72.45, pTM = 0.45).

**Supplementary Fig. 18.**
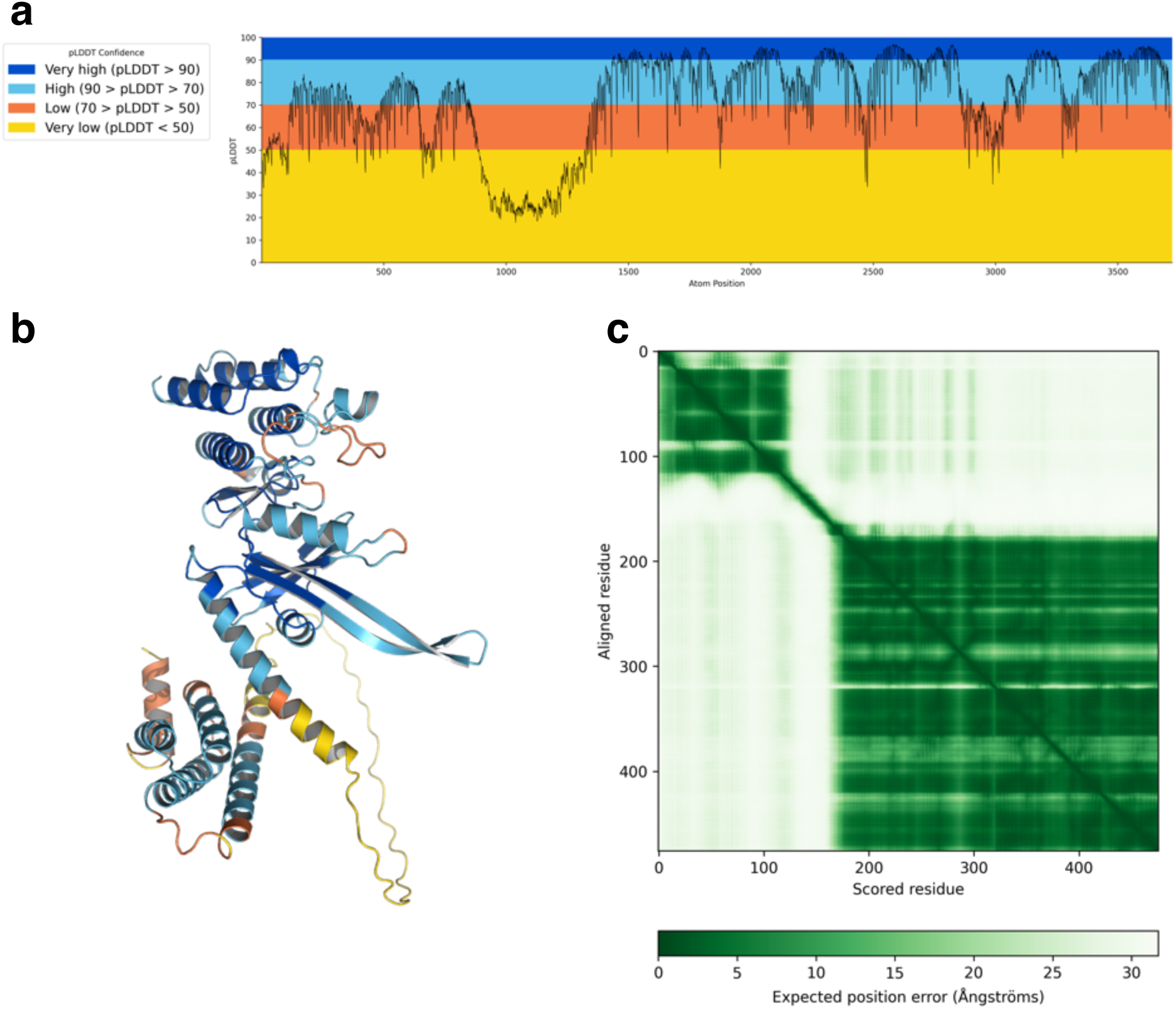
AlphaFold3 Prediction of Pm13. pLDDT plot (**a**), pLDDT coloured structure (**b**), and predicted aligned error plot (**c**) for the highest confidence prediction of Pm13 (Average pLDDT = 71.93, pTM = 0.62).

**Supplementary Fig. 19.**
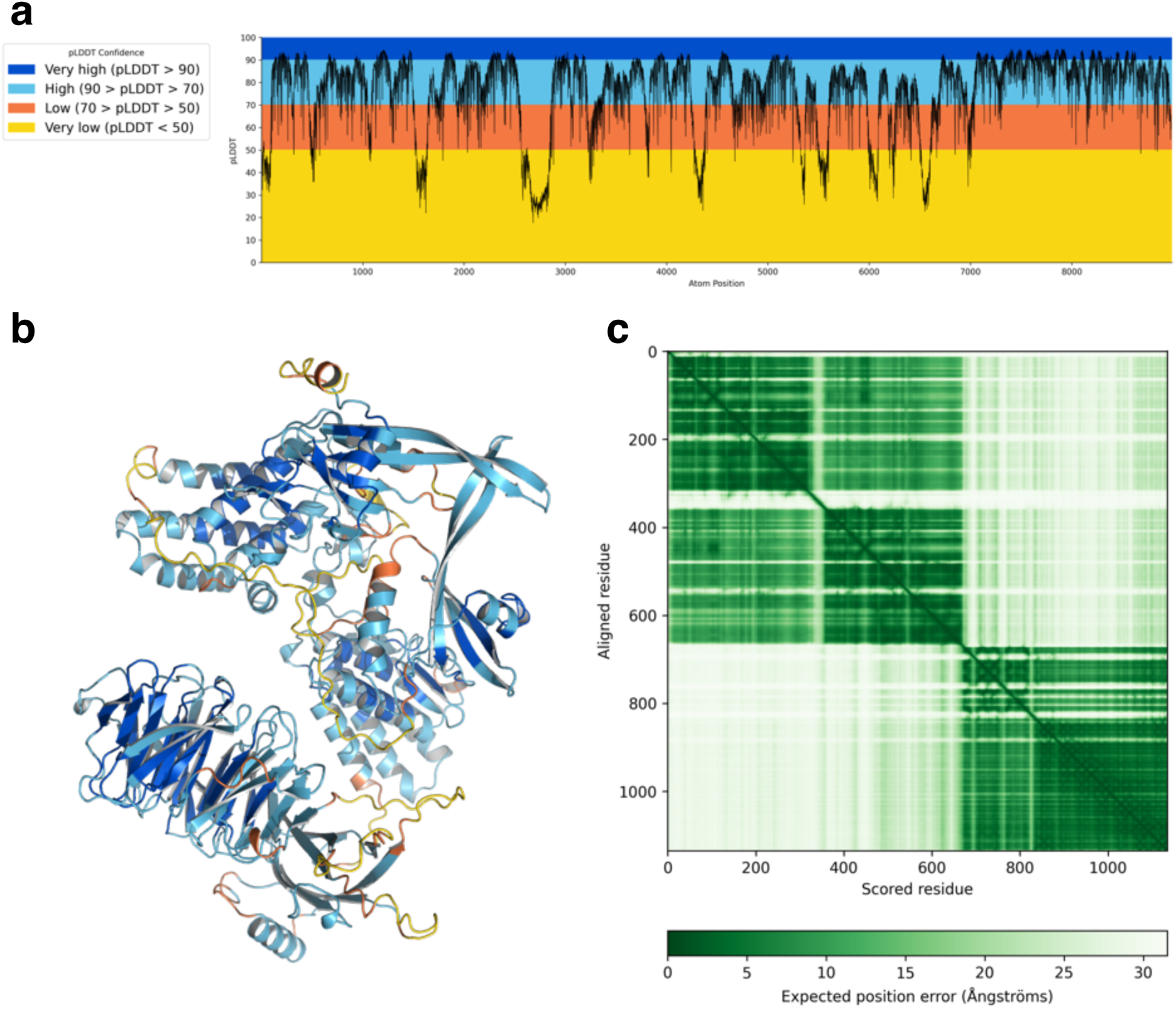
AlphaFold3 Prediction of Lr39. pLDDT plot (**a**), pLDDT coloured structure (**b**), and predicted aligned error plot (**c**) for the highest confidence prediction of Lr39 (Average pLDDT = 75.56, pTM = 0.57).

**Supplementary Fig. 20.**
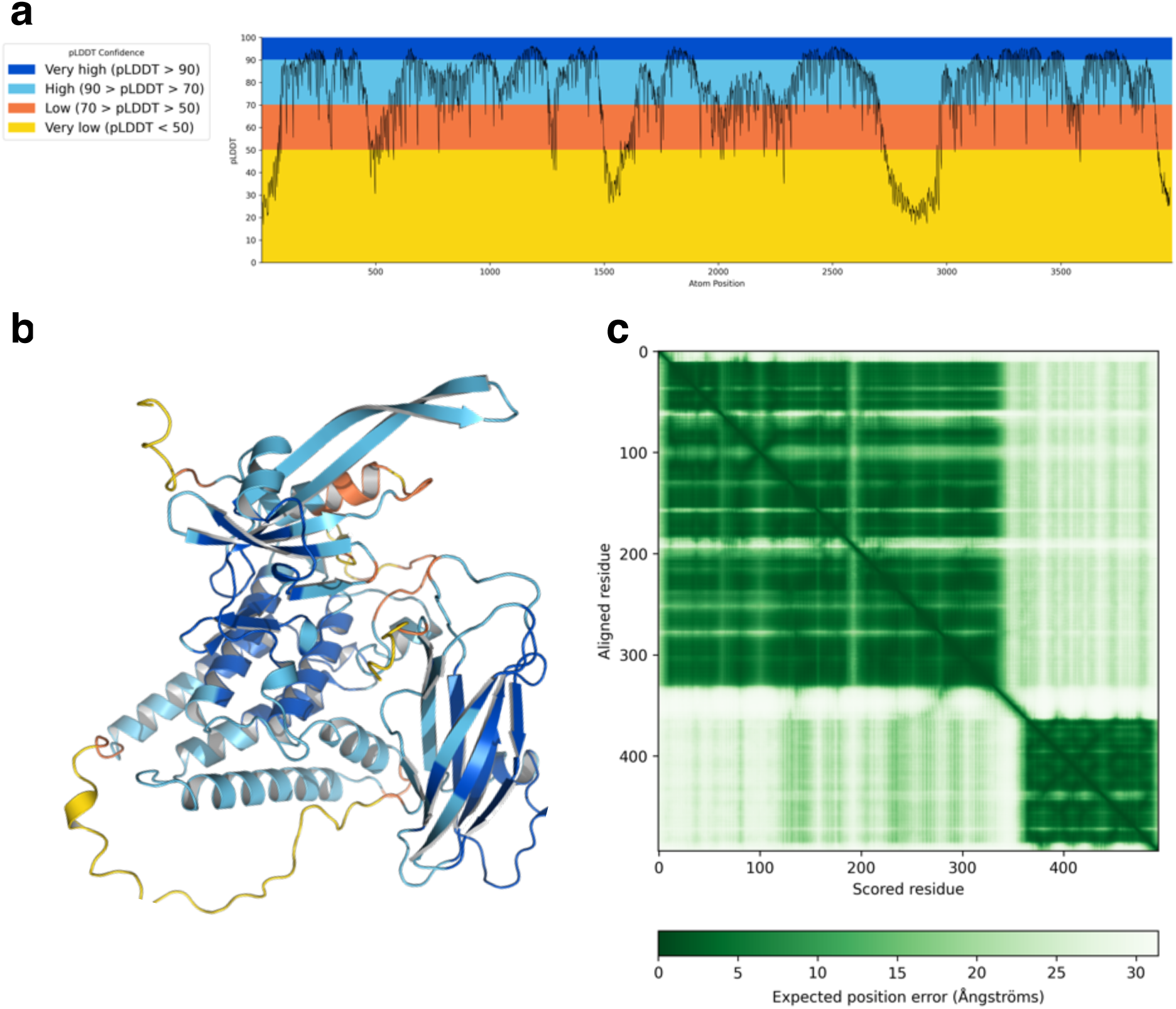
AlphaFold3 Prediction of Snn3-1B. pLDDT plot (**a**), pLDDT coloured structure (**b**), and predicted aligned error plot (**c**) for the highest confidence prediction of Snn3-1B (Average pLDDT = 75.19, pTM = 0.67).

**Supplementary Fig. 21.**
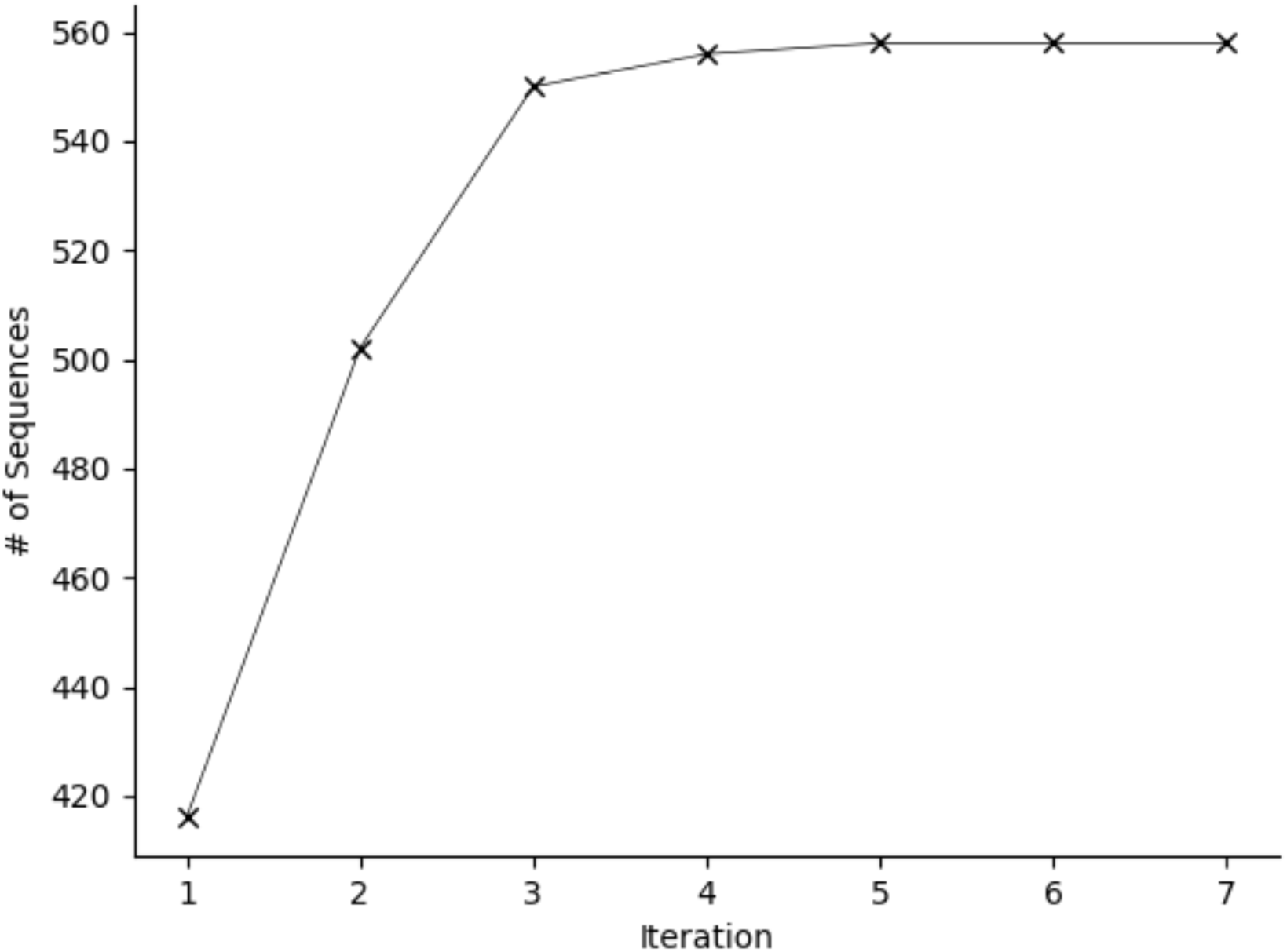
Iterative refinement of the extended β-finger HMM. The seed HMM of the extended β-finger motif, derived from the amino acid sequences of cloned KFPs, was iteratively refined against the Chinese Spring proteome until three consecutive iteration steps annotated the same number of sequences in the proteome (i.e., at step 7).

**Supplementary Fig. 22.**
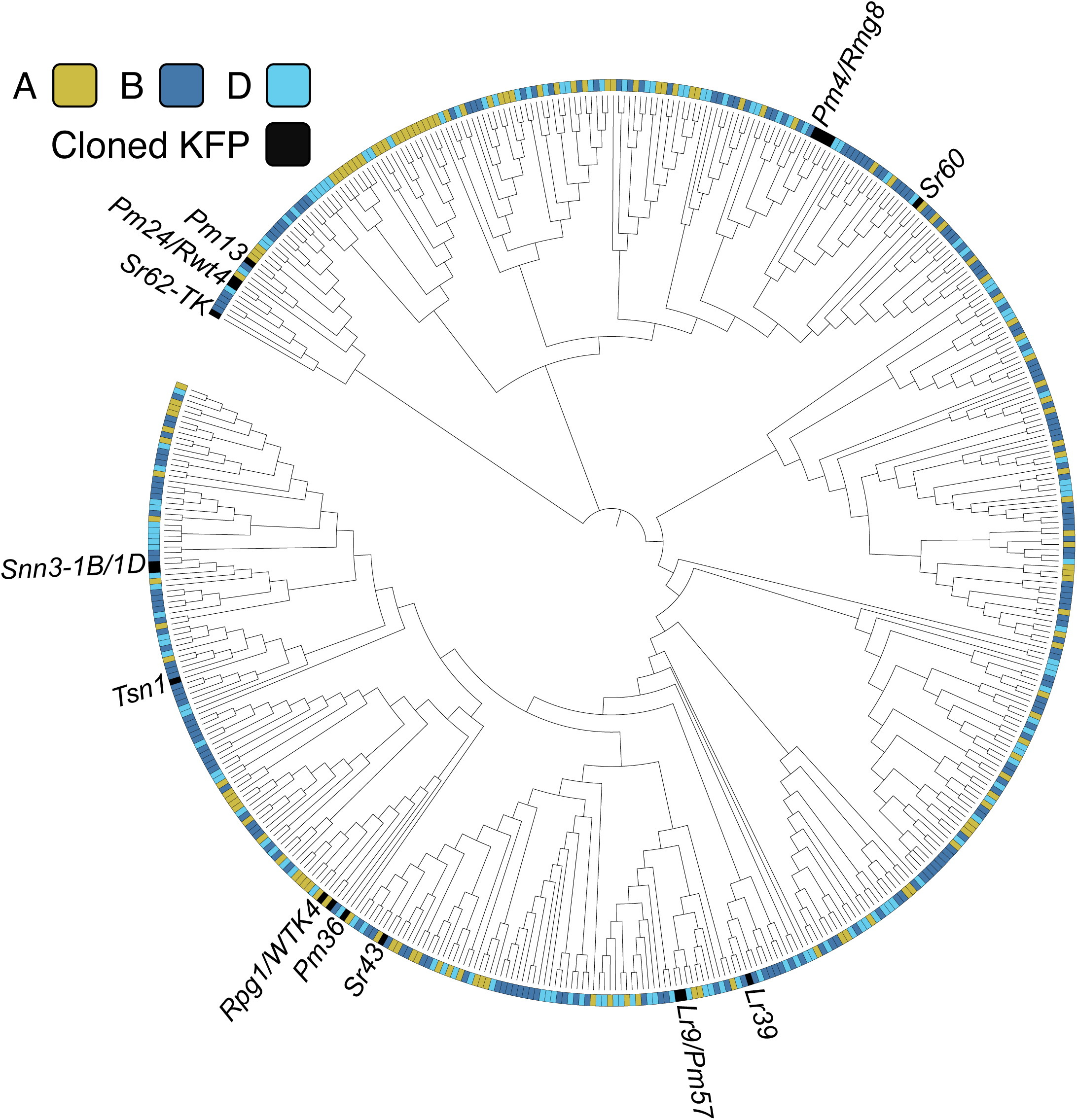
Distribution of KFP genes across the Chinese Spring subgenomes. Phylogenetic tree of KFP genes in the Chinese Spring genome. The subgenome origin of each KFP gene is colored according to the legend. Previously cloned KFP genes are shown in black and named.

**Supplementary Fig. 23.**
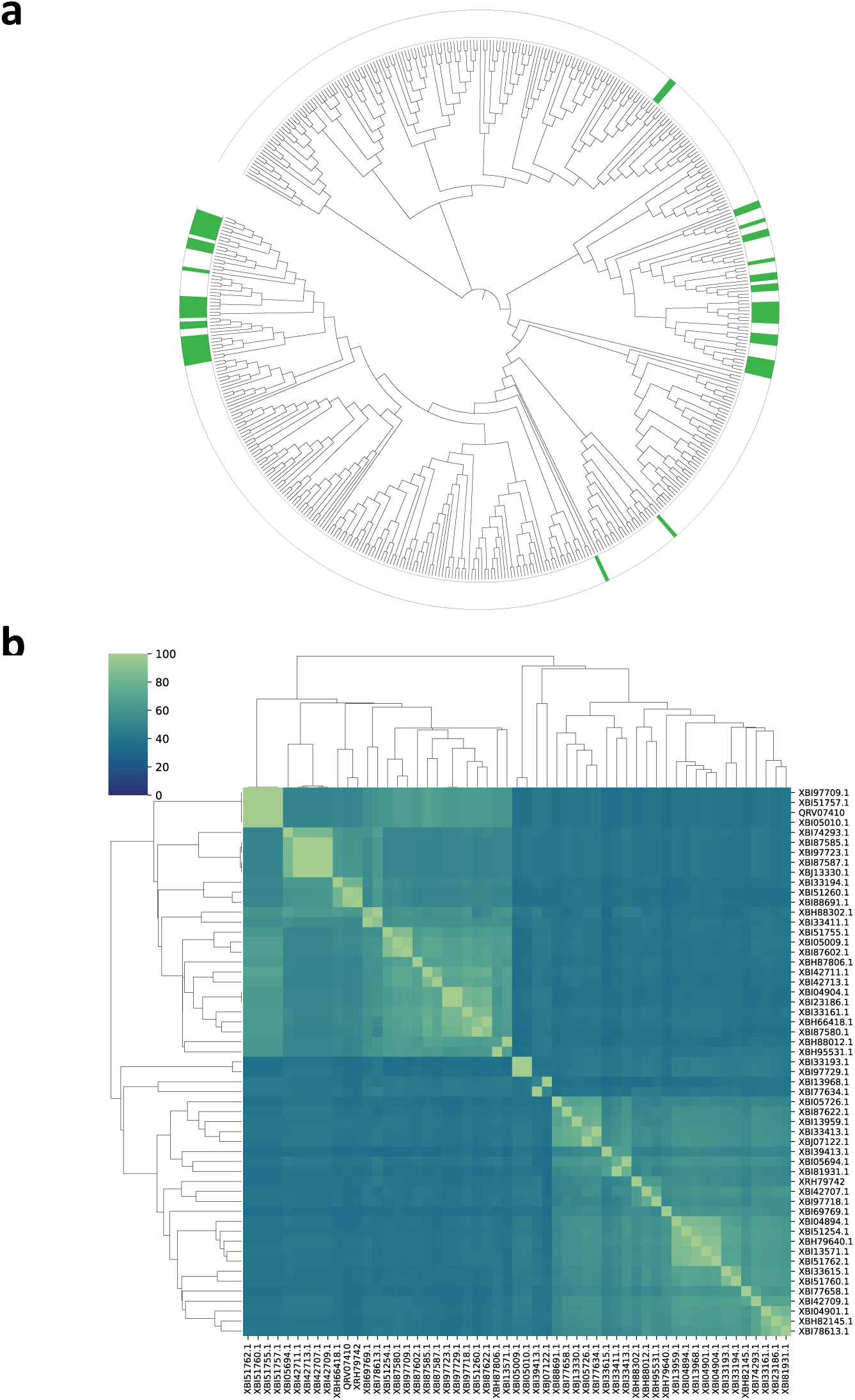
Distribution and similarity of KFPs with an MSP auxiliary module across the phylogenetic tree of KFPs from Chinese Spring. **a**, Distribution of MSP auxiliary modules across the phylogenetic tree of KFPs in Chinese Spring. KFPs with an MSP auxiliary module are shown in green. **b**, Percentage identity matrix of the extended β-finger kinase domains of KFPs with an MSP auxiliary module.

**Supplementary Fig. 24.**
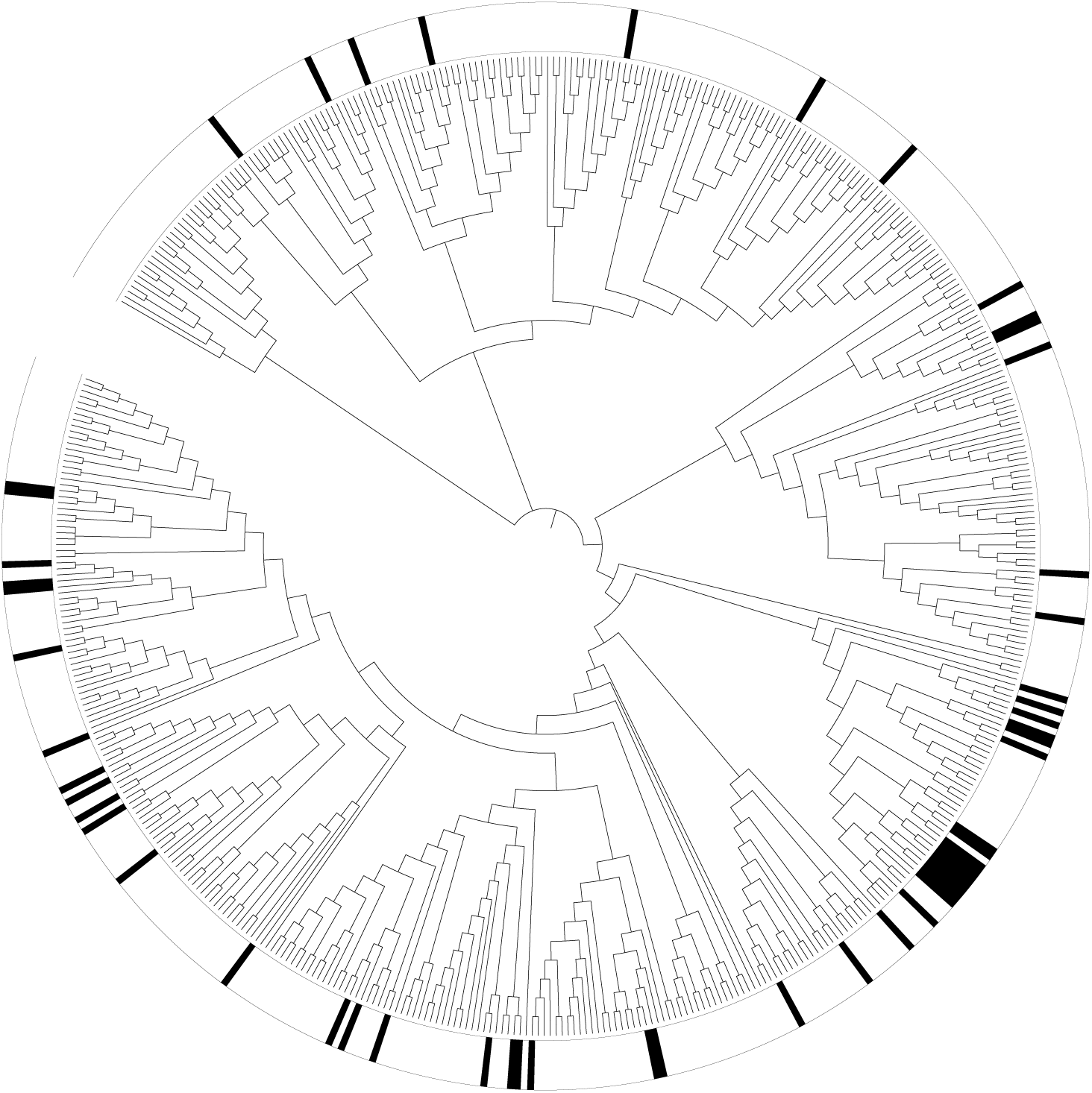
Distribution and similarity of KFPs with a kinase auxiliary module across the phylogenetic tree of KFPs from Chinese Spring. **a**, Distribution of kinase auxiliary modules across the phylogenetic tree of KFPs in Chinese Spring. KFPs with a kinase auxiliary module are shown in red. **b**, Percentage identity matrix of the extended b-finger kinase domains of KFPs with a kinase auxiliary module.

**Supplementary Fig. 25.**
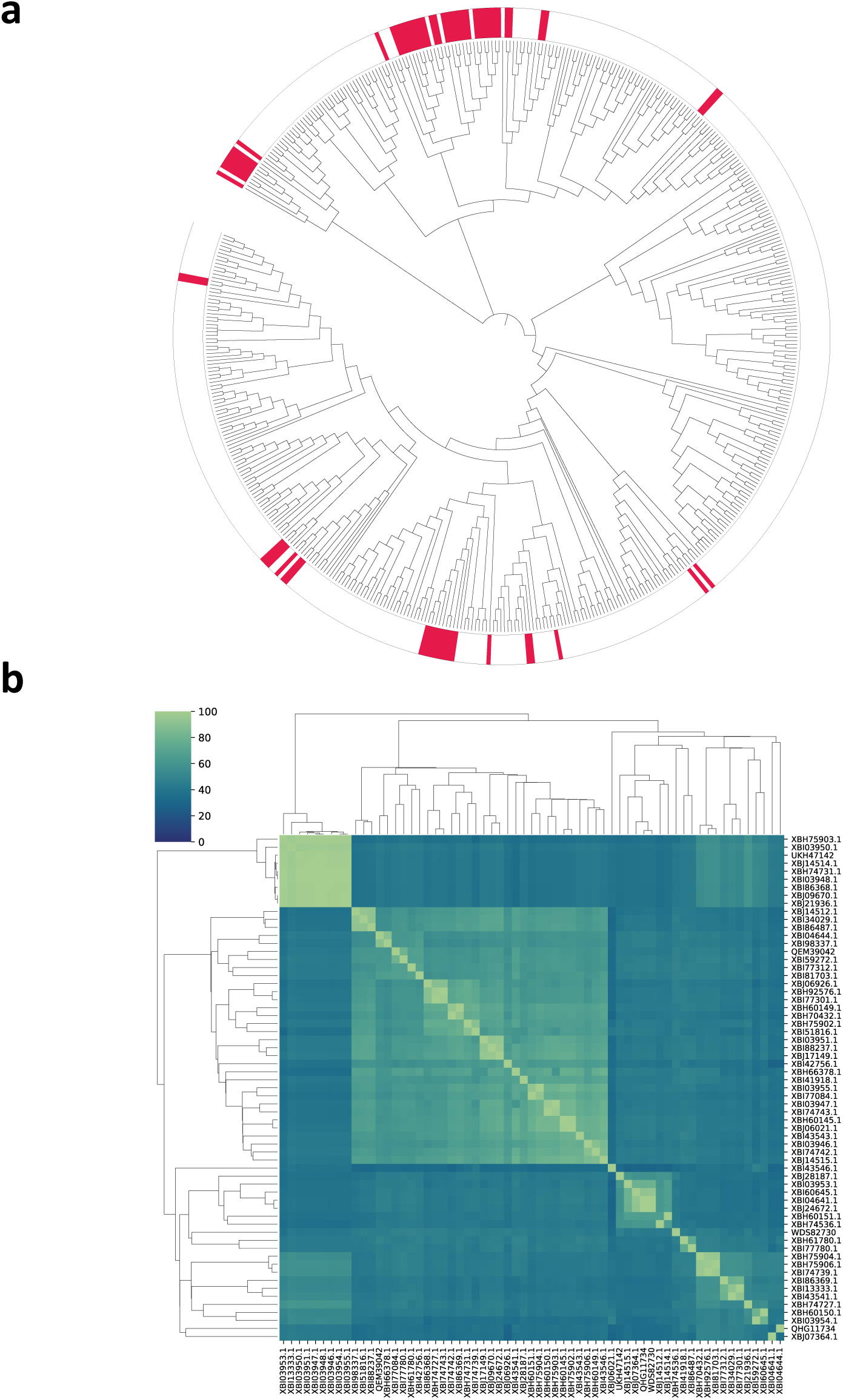
Distribution of KFPs with the extended β-finger but no auxiliary modules across the phylogenetic tree of KFPs in Chinese Spring. Phylogenetic tree of KFPs in Chinese Spring. The positions of extended β-finger kinases, *i.e.*, kinases with an extended β-finger but no fused auxiliary module, are shown in black.

**Supplementary Fig. 26.**
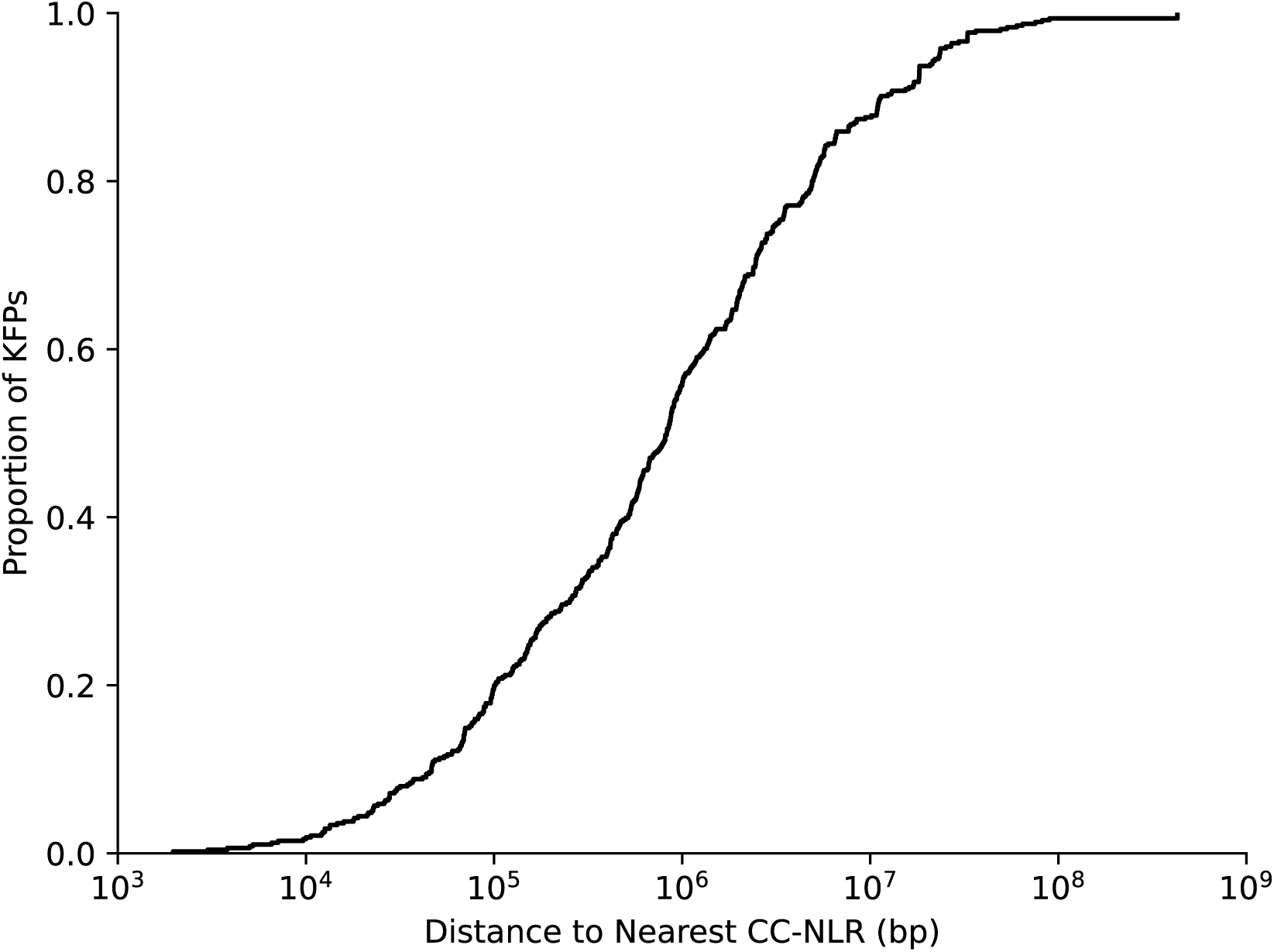
Distance to nearest CC-NLR gene for each KFP gene in the Chinese Spring genome. Cumulative distribution of the distance between each Chinese Spring KFP gene (n=476) and its nearest CC-NLR locus. About 28.2% of all KFP genes were located within 200 kb of a CC-NLR gene.

**Supplementary Fig. 27.**
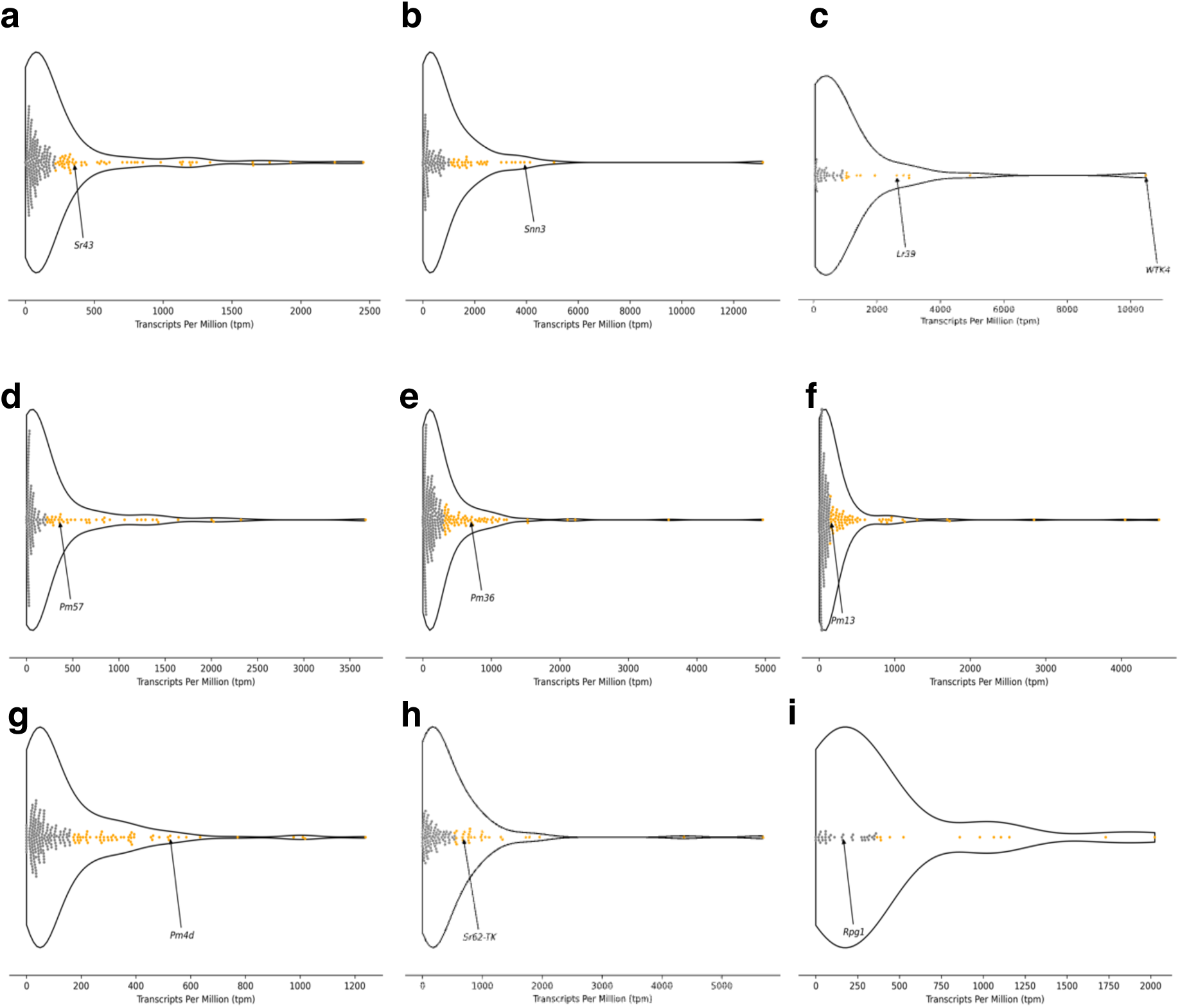
Functional KFP genes are among the most expressed KFP genes. Transcript abundance for KFP genes in *de novo* assembled transcriptomes of *Thinopyrum elongatum* (**a**), *Aegilops tauschii* (**b, c**), *Ae. searsii* (**d**), *T. turgidum* ssp. *dicoccoides* (**e**), *Ae. longissima* (**f**), *T. turgidum* (**g**), *Ae. sharonensis* (**h**), and *Hordeum vulgare* (**i**). Functionally validated KFP genes cloned from each species are highlighted. Transcript levels were estimated from self-aligned RNA-seq data as transcripts per million (TPM). The top 25% most highly expressed KFP genes are shown in orange.

**Supplementary Fig. 28.**
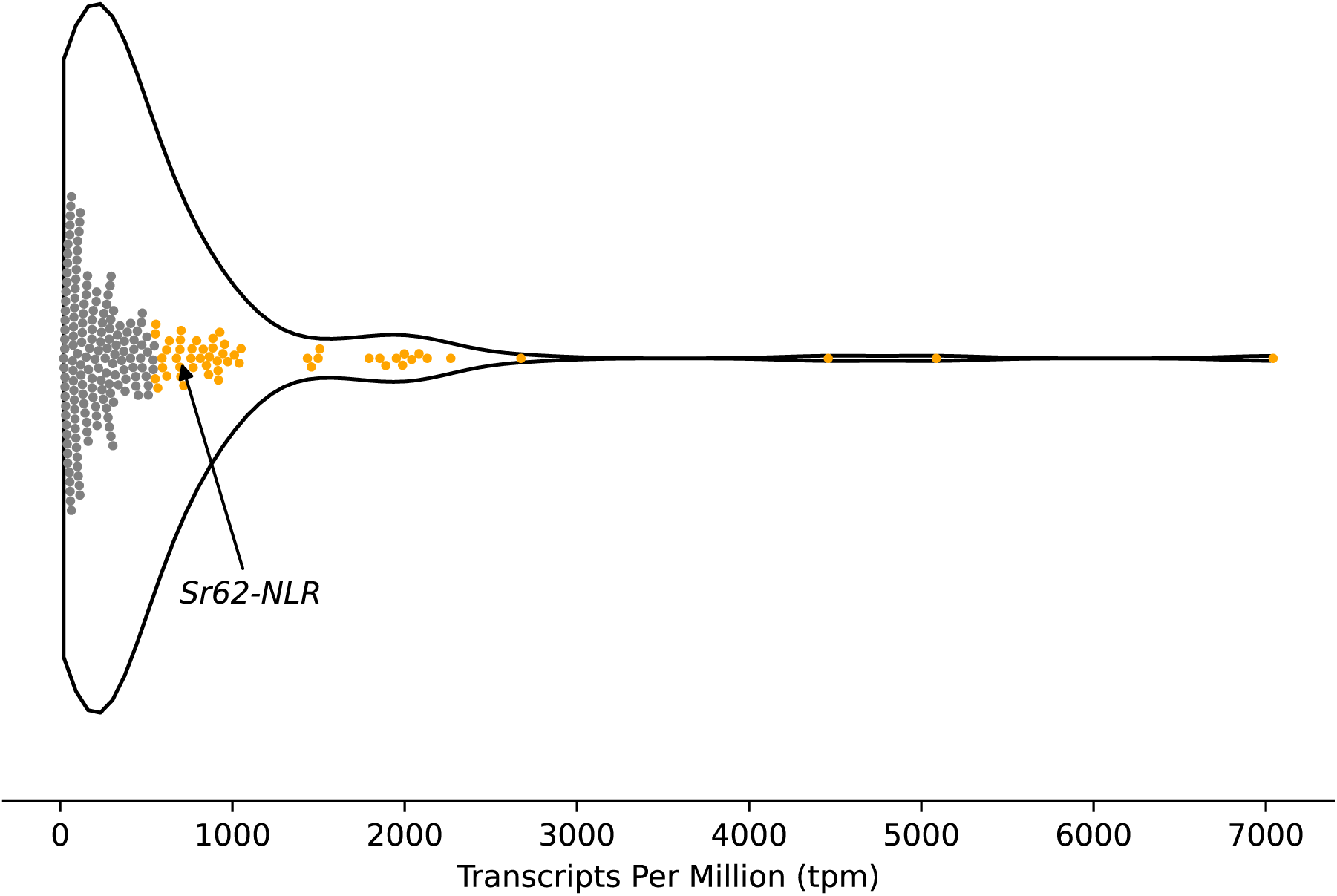
*Sr62^NLR^* is highly expressed among NLR genes. Transcript abundance of NLR genes in the *de novo* assembled transcriptome of *Aegilops sharonensis*. The functionally validated helper NLR gene of *Sr62^TK^*, *Sr62^NLR^*, is labeled. Transcript levels were estimated from self-aligned RNA-seq data as TPM. The top 25% most highly expressed NLR genes are shown in orange.

